# Mutant p53 Exploits Enhancers to Elevate Immunosuppressive Chemokine Expression and Impair Immune Checkpoint Inhibitors in Pancreatic Cancer

**DOI:** 10.1101/2024.08.28.609802

**Authors:** Dig B. Mahat, Heena Kumra, Sarah A. Castro, Emily Metcalf, Kim Nguyen, Ryo Morisue, William W. Ho, Ivy Chen, Brandon Sullivan, Leon K. Yim, Arundeep Singh, Jiayu Fu, Sean K. Waterton, Yu-Chi Cheng, Sylvie Roberge, Enrico Moiso, Vikash P. Chauhan, Hernandez Moura Silva, Stefani Spranger, Rakesh K. Jain, Phillip A. Sharp

## Abstract

Pancreatic ductal adenocarcinoma (PDAC) is an aggressive cancer without effective treatments. It is characterized by activating KRAS mutations and p53 alterations. However, how these mutations dysregulate cancer-cell-intrinsic gene programs to influence the immune landscape of the tumor microenvironment (TME) remains poorly understood. Here, we show that p53^R172H^ establishes an immunosuppressive TME, diminishes the efficacy of immune checkpoint inhibitors (ICIs), and enhances tumor growth. Our findings reveal that the upregulation of the immunosuppressive chemokine Cxcl1 mediates these pro-tumorigenic functions of p53^R172H^. Mechanistically, we show that p53^R172H^ associates with the distal enhancers of the Cxcl1 gene, increasing enhancer activity and Cxcl1 expression. p53^R172H^ occupies these enhancers in an NF-κB-pathway-dependent manner, suggesting NF-κB’s role in recruiting p53^R172H^ to the Cxcl1 enhancers. Our work uncovers how a common mutation in a tumor-suppressor transcription factor appropriates enhancers, stimulating chemokine expression and establishing an immunosuppressive TME that diminishes ICI efficacy in PDAC.

**Graphical Abstract:** 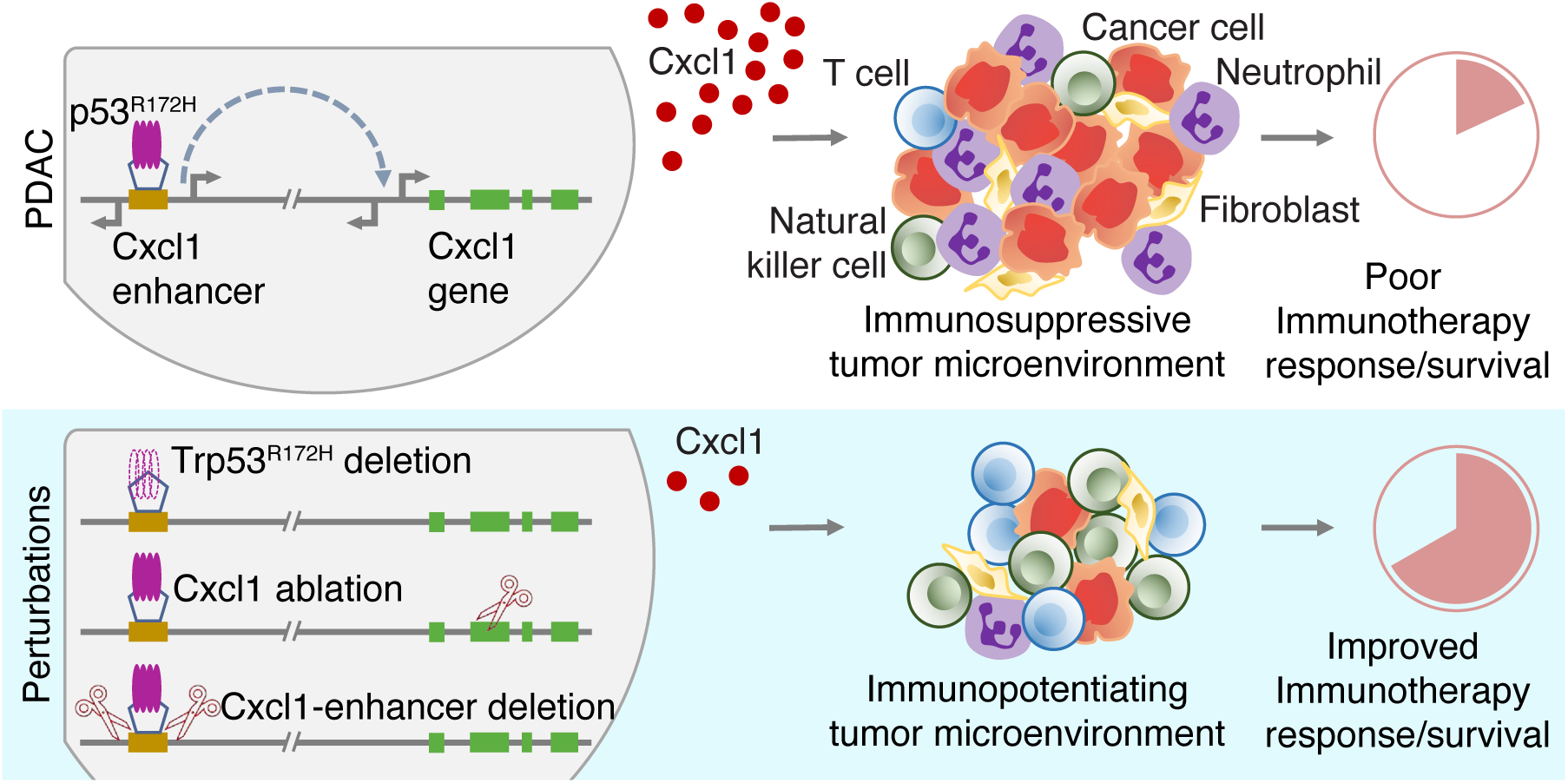

## Introduction

Pancreatic ductal adenocarcinoma (PDAC) is the third leading cause of cancer deaths due to late diagnosis and limited response to existing therapies^1^. It is predicted to become the second leading cause of cancer-related deaths by 2030^2^, highlighting the urgent need for new therapeutic strategies. One approach that could offer immediate benefits is enhancing PDAC’s susceptibility to existing therapeutic modalities, such as immune checkpoint inhibitors (ICIs). Despite their success in other cancers, ICIs have yet to show benefits for PDAC patients^3–7^. Recent strategies to harness the potential of ICIs by targeting additional factors such as DNA-repair proteins^8–10^, costimulatory receptors^11,12^, and immune modulators^13–15^ have shown promise in some cancers. Gaining deeper mechanistic insight into how oncogenic events shape cancer-cell-intrinsic gene programs to influence cell-extrinsic characteristics and sustain a pro-tumorigenic tumor microenvironment (TME) is crucial for unlocking this therapeutic approach.

About 90% of PDACs harbor activating KRAS mutations, suggesting it is a founding oncogenic event, and ∼70% have alterations in the TP53 tumor suppressor gene, suggesting the abrogation of the genome-guarding role of p53 accelerates the malignant progression^16^. The majority of p53 alterations are missense mutations in the DNA binding domain, which abolishes recognition of canonical binding motifs^17^. The six most frequently mutated p53 residues in human cancer are broadly categorized as ‘contact’ mutations (R248 and R273) or ‘structural’ mutations (R175, G245, R249, R282) based on defects in binding or structural deformation of the DNA-binding domain, respectively^18^. Mutant p53 exerts pro-tumorigenic effects either by loss-of-function^19,20^, a dominant-negative effect of suppressing the wild-type p53 function^21^, or by acquiring additional gain-of-function^22–24^. These mutant proteins can associate with transcription factors and other effectors in augmenting the transactivation potential of the interacting partners^25–33^. However, the interacting partners, affected pathways, and altered gene programs are highly context-dependent^34–38^.

The TME of PDAC is highly desmoplastic, hypoxic, and considered immunologically cold^39,40^. The low tumor mutational burden and the resulting low neo-antigen burden further insulate PDAC from the effectiveness of ICIs^41^. Consequently, understanding how cell-intrinsic programs shape the PDAC TME could reveal novel therapeutic opportunities. Over the past decade, some progress has been made^7^. For example, cancer-cell-specific Pin1 establishes a desmoplastic and immunosuppressive TME, and its pharmacological targeting appears to synergize with immunochemotherapy in the treatment of PDAC^42^. Similarly, mutant p53 harboring pancreatic ductal cells accumulate neutrophils and polymorphonuclear myeloid-derived suppressor cells that are anti-inflammatory and resist immunotherapy^43^. Still, the mechanistic insights on how mutant p53 contributes to immunosuppressive TME remain poorly understood.

We hypothesized that mutant p53 - which is highly stabilized compared to its wild-type counterpart^44,45^ and retains a functional and potent transactivation domain^46^ could drive new gene programs through mechanisms distinct from those of p53^WT^. For example, the transcription factor KLF5, which is frequently mutated in many cancers, gains over 5,000 new binding sites in comparison to wild-type KLF5, including super-enhancers proximal to genes that promote tumor growth^47^. Similarly, the splice variant of the androgen receptor that lacks the ligand binding domain occupies unique genomic sites and displays novel DNA binding motifs^48^. In this study, we examined the transcriptional influence of p53 missense mutations. We focused on the most prevalent p53 mutation^49^ (R175H in humans and R172H in mice) in the context of almost universally co-occurring Kras activating mutation (G12D) in PDAC. To investigate the role of p53^R172H^ in shaping the TME, we used cells derived from a genetically engineered mouse model (GEMM) of PDAC, created isogenic cell lines, and employed a host of techniques, including genome and transcriptome sequencing, CRISPR-Cas9 gene editing, immune profiling, and mouse survival studies with ICIs.

Our findings indicate that p53^R172H^ drives tumor progression by modulating cancer-cell-specific gene expression, which reprograms the TME into an immunosuppressive milieu. Specifically, p53^R172H^ is associated with lower T cell infiltration, higher myeloid-derived suppressor cell (MDSC) infiltration, and diminished efficacy of ICIs in mouse models of PDAC. At the molecular level, p53^R172H^ exerts its immunosuppressive effects primarily by regulating chemokine genes, particularly Cxcl1. p53^R172H^ associates with distal enhancers of Cxcl1 and amplifies its expression in an NF-κB-dependent manner. This interaction suggests a broader mechanism in which mutant p53 co-opts enhancers, which vastly outnumber gene promoters, through collaboration with other transcription factors. By interacting with the enhancer’s cognate binders, mutant p53 modulates cell-intrinsic gene expression that ultimately impacts the cell-extrinsic features of the TME. This study not only elucidates a mechanism of immune suppression driven by p53^R172H^ but also highlights broader implications for p53 missense mutations and their cooperation with other transcription factors in pancreatic cancer, offering potential targets for therapeutic intervention.

## Results

### p53^R172H^ regulates a subset of chemokines

To examine the transcriptional programs regulated by p53^R172H^, we used KPC cells derived from a GEMM of PDAC (*LSL-Kras^G12D/+^;LSL-Trp53^R172H/+^;Pdx-1-Cre*)^50^. Given the finding that more than 90% of cancer cases lose the *Trp53^WT^* allele in the presence of mutant p53 allele^51,52^, we used Exome-seq to check the status of *Trp53* alleles and confirmed that the *Trp53^WT^*allele is indeed lost (Figure S1A). This *Trp53^R172H/-^* genotype enables the examination of the gain-of-function effects of p53^R172H^, distinguishing it from the dominant-negative confounding effect in the parental *Trp53^R172H/+^* genotype.

To understand how p53^R172H^ alters gene programs, we compared transcriptional programs between *Trp53^R172H/-^* and *Trp53^-/-^*cells. However, instead of comparing with cells derived from other GEMMs of PDAC (such as *LSL-Kras^G12D/+^;LSL-Trp53^-/-^;Pdx-1-Cre*)^52–54^, which likely harbor non-overlapping mutations arising from the genomic instability due to a lack of p53^WT^ in both cases, we made isogenic *Trp53^-/-^* cells (*Kras^G12D/+^;Trp53^-/-^*) using CRISPR-Cas9 (Figure 1A) and selected single-cell clones and clonal mix population. Although all cells used in this study harbor *Kras^G12D/+^*, the genotypes hereafter are solely indicated by *Trp53* status for simplicity. We confirmed the deletion of Exon-2 to Exon-10 - identical region deleted to generate *Trp53^-/-^* mice^52^ - at the DNA, RNA, and protein levels (Figure S1B-D). We performed RNA-seq to measure the p53^R172H^-mediated gene expression changes in the Kras^G12D^ background and found several hundred genes under the control of p53^R172H^ (Figure 1B). Notably, the expression of canonical p53 target genes^55^ remained relatively unchanged, confirming the lack of wild-type p53 function in both cells. We found the genes upregulated by p53^R172H^ were enriched for chemokines and targets of the NF-κB transcription factor (Figure 1C). The genes suppressed by p53^R172H^ were enriched for negative regulation of biosynthesis and metabolism, likely related to the control of proliferation (Figure S1E). We performed a chemokine array to measure the levels of secreted chemokines in the tissue culture media and found a subset of chemokines differentially regulated by p53^R172H^ (Figure S1F). Most of the p53^R172H^-regulated chemokines belong to either CCL or ELR+CXCL chemokine subclass^56^.

**Figure 1.**
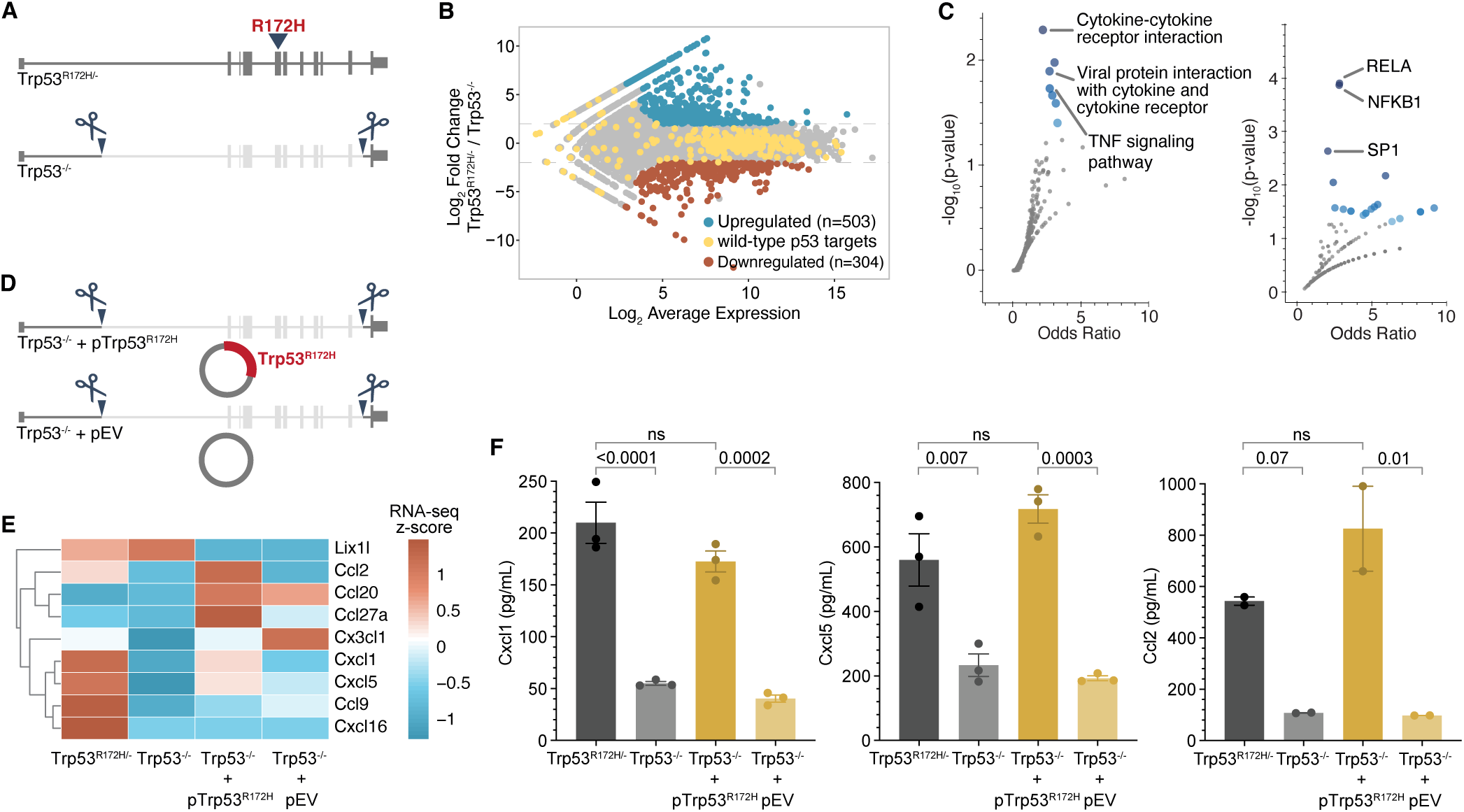
p53^R172H^ elevates the expression of a subset of chemokine genes. **A,** Generation of *Trp53^-/-^* isogenic cells from the parental *Trp53^R172H/-^*cells. *Trp53* gene with R172H mutation in exon-5 is deleted from intron-1 to intron-10 using CRISPR/Cas9. **B**, Minus-average (MA) plot of RNA-seq transcripts showing the differentially expressed genes between *Trp53^R172H/-^* and *Trp53^-/-^*isogenic cells. Significantly upregulated and downregulated genes (adjusted p-value < 0.001 and four-fold change in normalized counts) are shown in blue and red, respectively. The wild-type p53-regulated genes are shown in yellow. **C**, Pathways (left) and transcription factor targets (right) enriched in p53^R172H^-upregulated genes. Gene Ontology analysis was performed using Enrichr^1^^101^ against the KEGG pathway database^102^ (left) and TRRUST database^103^ (right). Blue dots represent significant gene sets (p-value < 0.05), and the darker color represents higher significance. **D**, Generation of *Trp53^R172H^*-restored isogenic cells in *Trp53^-/-^*cells using a *Trp53^R172H^* cDNA expression cassette in piggyback vector (*Trp53^-/^*^-^ + *pTrp53^R172H^*). An empty vector without the *Trp53^R172H^* cDNA expression cassette (*Trp53^-/-^* + *pEV*) was inserted in *Trp53^-/^*^-^ as a control. **E**, mRNA levels of the expressed chemokine genes are shown as the z-score heatmap of RNA- seq transcripts per million (TPM) in the four isogenic cells. **F**, Quantification of the three chemokine genes under p53^R172H^ control by ELISA in the tissue culture media of the four isogenic cells. P-values are calculated from a one-way ANOVA test followed by a post hoc test with Benjamini-Hochberg correction. Panels B, C, E, and F use *Trp53^-/-^* clone-1 isogenic cells.

To exclude clonal variability as a confounding factor, we performed RNA-seq experiments with an additional single-cell clone of isogenic *Trp53^-/-^*cells, as well as a clonal-mix population generated by pooling three single-cell clones. Our results showed a similar number of p53^R172H^- regulated genes and the dependence of a similar set of chemokine genes on p53^R172H^ across clonal and clonal-mix populations (Figure S2A). The genes upregulated and downregulated upon the deletion of *Trp53^R172H^* highly overlapped among clonal and clonal-mix populations (Figure S2B), and the overlapping genes showed enrichment for chemokine genes (Figure S2C), as observed previously with a representative clone. Similarly, enhanced secretion of Cxcl1, Cxcl5, and Ccl2 proteins was dependent on p53^R172H^ in the clonal-mix population as measured by ELISA in tissue-culture media (Figure S2D).

We further substantiated the generality of p53^R172H^-mediated transcriptional regulation and its influence on chemokine gene expression using syngeneic *Trp53^-/-^* cells^57^, generated from a GEMM of PDAC (*p48-Cre;Kras^LSL-G12D/+^;Trp53^loxP/+^*)^58^. Similar to the parental KPC cells, we found that the *Trp53^WT^* allele is lost in these PDAC cells (Figure S3A), likely due to the loss-of-heterozygosity, which we confirmed at the mRNA and protein levels (Figure S3B-C). RNA-seq analysis of the syngeneic *Trp53^-/-^*cells, compared with the *Trp53^R172H/-^* cells, highlighted the dependency of similar chemokine genes on p53^R172H^ (Figure S3D-E). Notably, specific chemokines—Cxcl1, Cxcl5, and Ccl2—showed reduced secretion, as measured by cytokine array (Figure S3F).

To determine whether p53^R172H^ is sufficient to regulate chemokine expression, we ectopically expressed *Trp53^R172H^*cDNA in *Trp53^-/-^* using the PiggyBac expression vector (Figure 1D, see Methods). An empty vector (pEV) served as a control. We confirmed the expression of p53^R172H^ (Figure S1G) and observed that mRNA levels of certain chemokine genes were partially restored in p53^R172H^-expressing *Trp53^-/-^* cells (Figure 1E). Further validation using ELISA confirmed that the reintroduction of p53^R172H^ in *Trp53^-/-^* cells significantly restored the expression of specific chemokines—Cxcl1, Cxcl5, and Ccl2 (Figure 1F). Collectively, these findings show that p53^R172H^ acquires a novel transcriptional regulatory function, driving the expression of a specific set of chemokine genes.

### p53^R172H^ creates an immunosuppressive TME and abrogates ICI efficacy

To test whether the loss of *Trp53^R172H^* suppresses the PDAC formation and growth, we orthotopically implanted *Trp53^R172H/-^* or *Trp53^-/-^* cells in the pancreas of immunocompetent wild-type mice. We profiled the immune landscape of the PDAC TME 21 days after implantation (Figure 2A). The tumors formed by *Trp53^R172H/-^*cells were significantly larger compared to the *Trp53^-/-^* cells (Figure 2B), indicating the role of p53^R172H^ in PDAC formation and growth. Immune profiling of the tumors using fluorescence-activated cell sorting (FACS) showed that the *Trp53^-/-^*tumors had higher infiltration of T cells (CD45^+^CD3e^+^CD8^+^), activated T cells (CD45^+^CD3e^+^CD8^+^CD44^+^), cytotoxic T cells (CD45^+^CD3e^+^CD8^+^Gzmb^+^) and activated conventional CD4^+^ T cells (CD45^+^CD3e^+^CD8^-^CD4^+^CD44^+^Foxp3^-^) compared to the *Trp53^R172H/-^* tumors (Figure 2C). In contrast, *Trp53^-/-^* tumors had lower infiltration of MDSCs (CD45^+^CD11b^+^Gr-1^+^Arg-1^+^) than *Trp53^R172H/-^* tumors (Figure 2D). The tumors established with orthotopic implantation of the clonal-mix population of *Trp53^-/-^* cells were also smaller and showed higher infiltration of T cells, activated T cells, cytotoxic T cells, and activated conventional CD4^+^ T cells (Figure S4A-B). We used immunofluorescence to further examine the immune composition of the tumors. We found a higher number of cytotoxic T cells (CD8^+^) and a lower number of neutrophils (Gr-1^+^) and MDSCs (Gr-1^+^Arg-1^+^) in *Trp53^-/-^* cells (Figure 2E).

**Figure 2.**
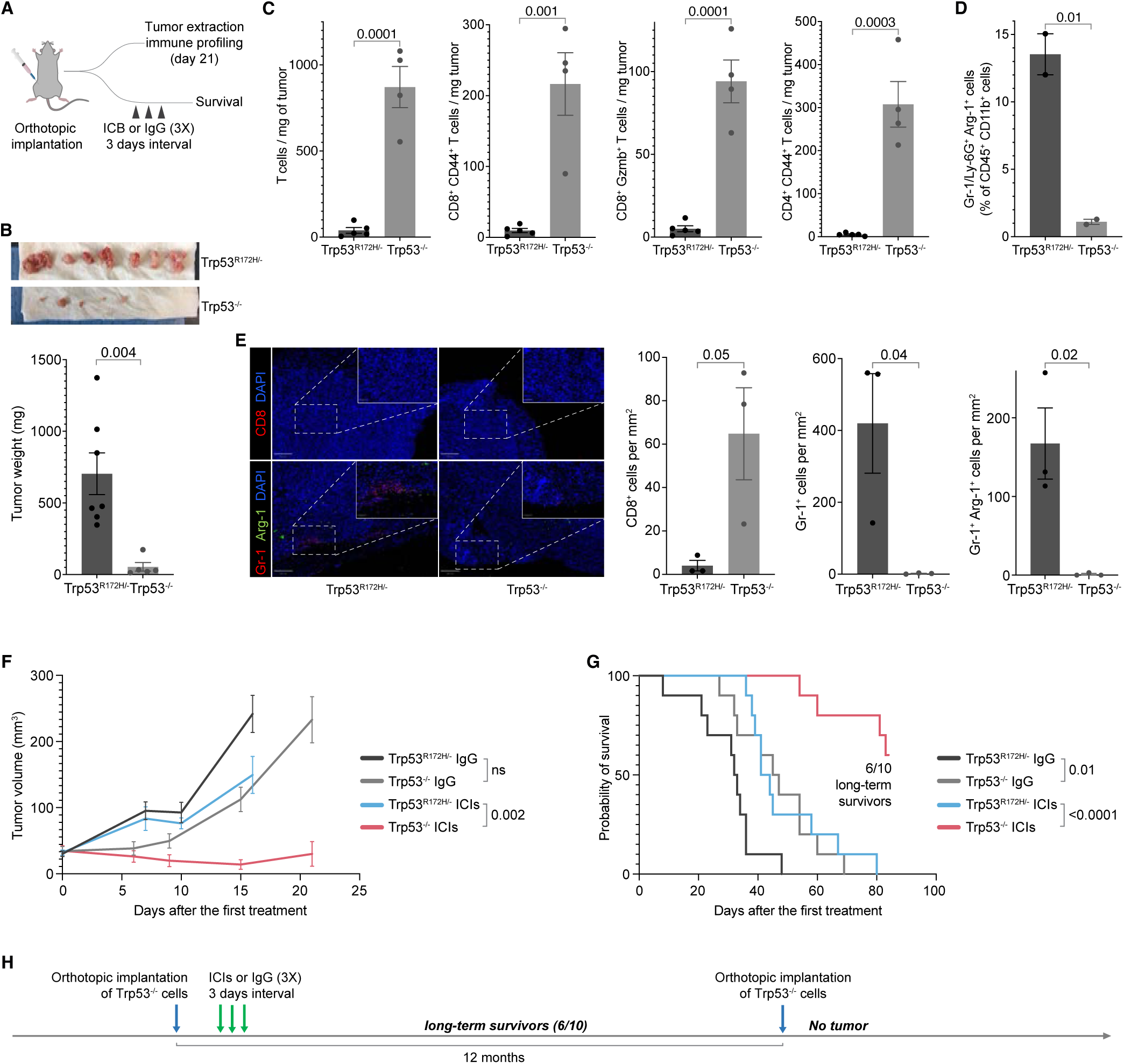
p53^R172H^ creates immunosuppressive TME and abrogates ICIs efficacy. **A,** Schematics and the experimental design of *Trp53^R172H/-^* and *Trp53^-/-^* isogenic cells’ orthotopic implantation in mouse pancreas, immune profiling of the tumors, ICIs treatment, and survival analysis. **B**, Weights of the *Trp53^R172H/-^* and *Trp53^-/-^* tumors. The p-value is calculated using a two-tailed t-test. **C**, Effect of the *Trp53* status on T cell infiltration in PDAC tumors. The p-values are calculated using a two-tailed t-test. **D**, Effect of the *Trp53* status on MDSCs infiltration in PDAC tumors. The p-value is calculated using a two-tailed t-test. **E**, Representative multiplex immunofluorescence staining of *Trp53^R172H/-^*and *Trp53^-/-^* tumors (left), and quantification of immune cells density from randomly selected 500 x 500 µm regions (right). The scale bars correspond to 200 μm and 50 μm in the main view and the magnified view, respectively. P-values are calculated using a two-tailed t-test. **F**, Effect of the *Trp53* status and ICIs on PDAC tumor growth. Control mice were treated with IgG. Tumor volumes from the last measurement with at least three mice left in the cohort were used to calculate p-values using a two-tailed t-test. **G**, Kaplan-Meier survival curves showing the effect of *Trp53* status and ICIs on the survival of mice implanted with either *Trp53^R172H/-^* or *Trp53^-/-^* cells. Control mice were treated with IgG. P-values are calculated using a log-rank (Mantel-Cox) test. **H,** Experimental design and timeline of tumor challenge experiment in the long-term survivor mice implanted with *Trp53^-/-^* PDAC tumors and treated with ICIs.

We then tested whether the increased CD8^+^ T cell infiltration in the TME of *Trp53^-/-^* tumors compared to *Trp53^R172H/-^* tumors would sensitize the tumors to ICIs. We treated the *Trp53^R172H/-^*or *Trp53^-/-^* cells-implanted mice with ICIs (anti-CTLA4 + anti-PD-1 combinatorial therapy) or IgG (control) three times in 3-day intervals and monitored the tumor growth and survival (Figure 2A). We used ultrasound to monitor the growth of PDAC tumors after ICIs administration until the mice reached a humane endpoint. In multiple replicate studies, we consistently observed a slower growth of *Trp53^-/-^* tumors than the *Trp53^R172H/-^* tumors (Figure 2F & Figure S4C). More importantly, the ICIs treatment dramatically increased the survival of mice implanted with *Trp53^-/-^* tumors (Figure 2G & Figure S4D). The ICIs treatment resulted in complete regression of *Trp53^-/-^* tumors in 6 out of 10 in the first cohort and 4 out of 8 in the second cohort, and these mice survived long-term. We challenged these long-term survivors with another orthotopic implantation of *Trp53^-/-^* cells a year after the first implantation and treatment. These mice were able to prevent tumor formation (Figure 2H), suggesting the establishment of immune memory. These *in vivo* studies indicate a role for p53^R172H^ in establishing an immunosuppressive TME and facilitating evasion of anti-tumor immunity in PDAC.

### Cxcl1 mediates an oncogenic role of p53^R172H^

We hypothesized that p53^R172H^ establishes an immunosuppressive TME by regulating the expression of immune-modulating genes. To investigate this, we focused on p53^R172H^- dependent chemokines, particularly Ccl2, due to its strong association with p53^R172H^ and its potential as a therapeutic target in PDAC^59^. Ccl2 is known to attract monocytes^60–62^ and promotes infiltration of natural killer (NK) cells^63,64^. Using CRISPR-Cas9, we generated isogenic *Ccl2^-/-^* cells while retaining *Kras^G12D/+^* and *Trp53^R172H/-^* mutations (Figure S5A). The frameshift mutations in the *Ccl2* gene resulted in the complete loss of Ccl2 expression (Figure S5B). Surprisingly, the absence of Ccl2 did not affect tumor growth upon orthotopic implantation (Figure S5C), nor did it enhance responsiveness to ICIs, unlike what was observed in *Trp53^-/-^* cells (Figure S5D). These findings led us to explore other p53^R172H^-dependent chemokines. We deleted the *Cxcl1* or *Cxcl5* gene in the *Ccl2^-/-^* cells and implanted the double knockout cells (*Ccl2^-/-^;Cxcl1^-/-^*or *Ccl2^-/-^;Cxcl5^-/-^*) in the pancreas of immunocompetent mice. The double knockout cells resulted in significantly smaller tumors (Figure S5E).

These results raised the question of whether the deletion of Cxcl1 or Cxcl5 alone could have impeded tumor growth. To address this, we made isogenic *Cxcl1^-/-^* or *Cxcl5^-/-^* cells using CRISPR-Cas9 while retaining the *Kras^G12D/+^* and *Trp53^R172H/-^*mutations (Figure 3A). The frameshift mutation resulted in a complete loss of the Cxcl1 expression (Figure S5F). Orthotopic implantation of *Trp53^R172H/-^;Cxcl1^-/-^* or *Trp53^R172H/-^;Cxcl5^-/-^* cells in the pancreas of wild-type mice demonstrated that the loss of Cxcl1 significantly reduced tumor size, recapitulating the effects of p53^R172H^ loss (Figure 3B). In contrast, Cxcl5 deletion alone did not impact tumor growth. Examination of the immune landscape of *Cxcl1^-/-^* tumors revealed a higher infiltration of T cells, activated T cells, cytotoxic T cells, and activated conventional CD4^+^ T cells, similar to the *Trp53^-/-^* tumors (Figure 3C). These results indicate that Cxcl1 contributes to p53^R172H^-mediated immunosuppression.

**Figure 3.**
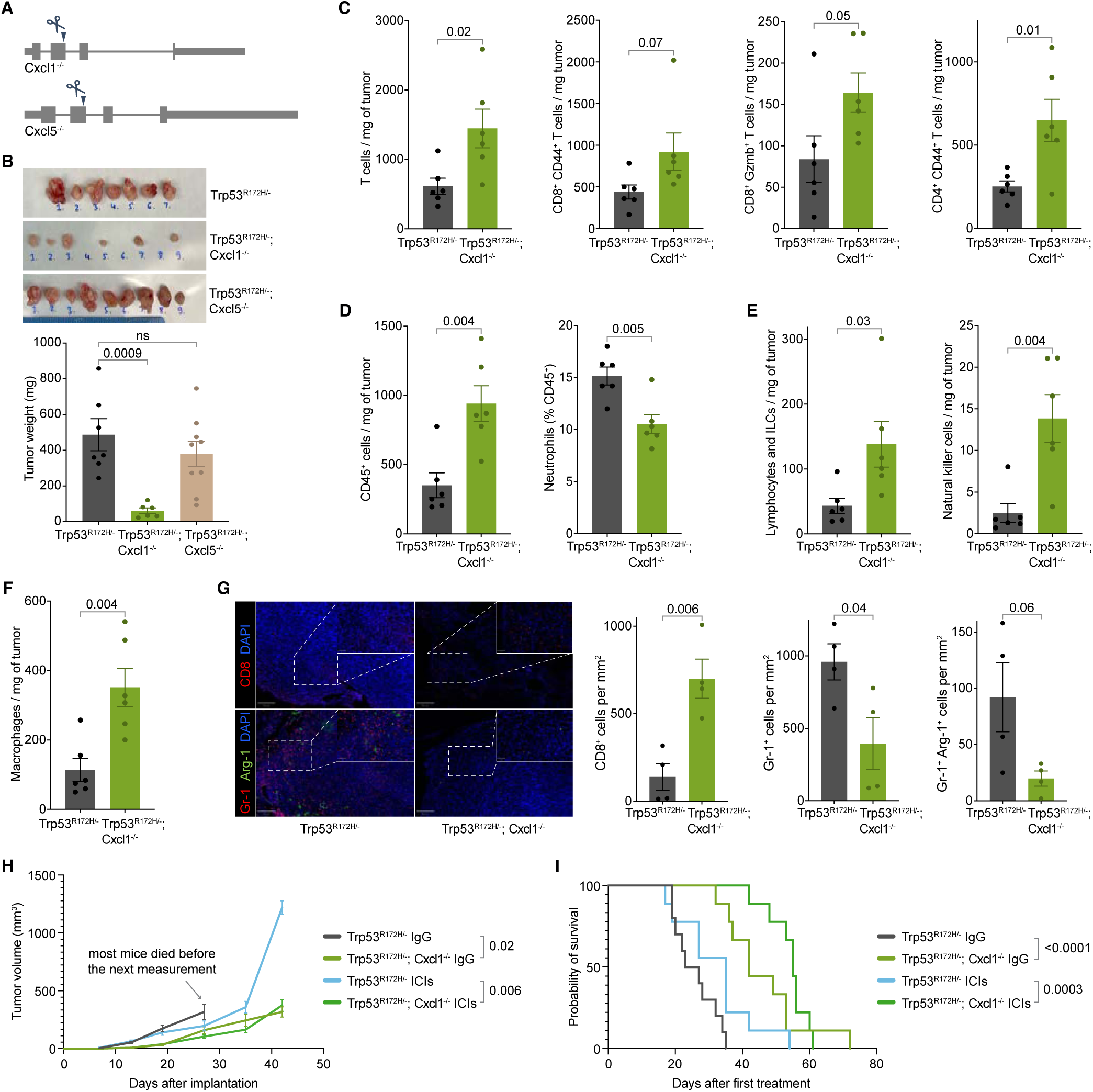
Cxcl1 mediates the immunosuppressive role of p53^R172H^ and abrogates ICIs efficacy. **A,** Generation of *Trp53^R172H/-^*;*Cxcl1^-/-^* and *Trp53^R172H/-^*;*Cxcl5^-/-^*isogenic cells from the parental *Trp53^R172H/-^* cells. A single guide-RNA-mediated genome editing using CRISPR/Cas9 resulted in a frameshift mutation. **B**, Weights of the *Trp53^R172H/-^*;*Cxcl1^-/-^* and *Trp53^R172H/-^*;*Cxcl5^-/-^*tumors compared with the *Trp53^R172H/-^* tumors. P-values are calculated from a one-way ANOVA test followed by a post hoc test with Benjamini-Hochberg correction. **C**, Effect of the *Cxcl1* status in T cell infiltration in PDAC tumors. P-values are calculated using a two-tailed t-test. **D**, Effect of the *Cxcl1* status in leukocyte (CD45^+^) and neutrophil (CD45^+^CD11b^+^MHCII^-^Ly-6G^+^) infiltration in PDAC tumors. P-values are calculated using a two-tailed t-test. **E**, Effect of the *Cxcl1* status in lymphocyte (CD45^+^CD11b^-^CD90^+^NK1.1^-^) and NK cell (CD45^+^CD11b^-^CD90^+^NK1.1^+^) infiltration in PDAC tumors. P-values are calculated using a two-tailed t-test. **F,** Effect of the *Cxcl1* status in macrophage (CD45^+^CD11b^+^CD64^+^) infiltration in PDAC tumors. P-values are calculated using a two-tailed t-test. **G,** Representative multiplex immunofluorescence staining of *Trp53^R172H/-^* and *Trp53^R172H/-^;Cxcl1^-/-^* tumors (left), and quantification of immune cells density from randomly selected 500 x 500 µm regions (right). The scale bars correspond to 200 μm and 50 μm in the main view and the magnified view, respectively. P-values are calculated using a two-tailed t-test. **H,** Effect of the *Cxcl1* status and ICIs in PDAC tumor growth. Control mice were treated with IgG. *Trp53^R172H/-^* tumor growth cohort is the same as in Figure S4C. Tumor volumes from the last measurement with at least three mice left in the cohort were used to calculate p-values using a two-tailed t-test. **I**, Kaplan-Meier survival curves showing the effect of *Cxcl1* status and ICIs in the survival of mice implanted with parental *Trp53^R172H/-^* or *Trp53^R172H/-^;Cxcl1^-/-^* cells. Control mice were treated with IgG. *Trp53^R172H/-^* mice survival cohort is the same as in Figure S4D. P-values are calculated using a log-rank (Mantel-Cox) test.

Cxcl1 is a chemotactic cytokine with a widely reported role in cancer^65^. The primary receptor for Cxcl1 is Cxcr2, which is mainly expressed in neutrophils^66^. Neutrophils can be recruited to the TME and differentiate into polymorphonuclear MDSCs (PMN-MDSCs), which contribute to a pro- tumorigenic environment^67^. We observed a reduced abundance of neutrophils among total CD45^+^ cells in *Trp53^R172H/-^;Cxcl1^-/-^*tumors (Figure 3D). We investigated other immune cell counterparts that could contribute to slower tumor growth. We observed an increase in infiltration of lymphocytes and innate lymphoid cells (CD45^+^CD11b^-^CD90^+^NK1.1^-^), natural killer (NK) cells (CD45^+^CD11b^-^CD90^+^NK1.1^+^), and macrophages (CD11b^+^CD64^+^) upon the loss of Cxcl1 (Figure 3E & Figure 3F), all of which have been reported to show anti-tumor activity^68^. Finally, we observed that the lymphocyte-to-neutrophil and NK-cell-to-neutrophil ratio was significantly higher in Cxcl1-ablated tumors, which suggests a TME with more robust anti-tumor activity (Figure S5G). We used immunofluorescence to further examine the immune composition of the tumors. We found a higher number of cytotoxic T cells (CD8^+^) and a lower number of immunosuppressive neutrophils (Gr-1^+^) and MDSCs (Gr-1^+^Arg-1^+^) in *Trp53^R172H/-^;Cxcl1^-/-^* tumors (Figure 3G).

We then tested whether the immune state of the TME in *Trp53^R172H/-^;Cxcl1^-/-^*tumors impedes tumor growth and elicits a response to ICIs. We found that Cxcl1 ablation slowed tumor growth (Figure 3H). More importantly, mice with *Trp53^R172H/-^;Cxcl1^-/-^* tumors exhibited better survival rates, with ICI treatment further improving survival (Figure 3I). Overall, our data suggest that one of the pro-tumorigenic pathways exploited by p53^R172H^ involves the elevation of Cxcl1 expression, which contributes to the establishment of an immunosuppressive TME, promoting tumor growth and diminishing the efficacy of ICIs.

### p53^R172H^ occupies Cxcl1 enhancers and increases their transcription activation potential

The significance of Cxcl1 in shaping the PDAC TME prompted us to investigate the mechanism by which p53^R172H^ regulates Cxcl1. p53 is a potent transcription factor with a robust transactivation domain, located in the N-terminus of the protein, that remains largely unaffected in mutant p53 proteins found in cancers. Most p53 mutations in cancers occur within the DNA-binding domain, resulting in the loss of the ability to bind the canonical DNA motifs recognized by wild-type p53.

Given its transcriptional activation potential, we investigated whether p53^R172H^ associates with new genomic sites. After testing nine different antibodies, we identified two that performed well with CUT&RUN^69^. In *Trp53^R172H/-^* cells, we observed significant p53^R172H^ occupancy at the Cxcl1 promoter (Figure 4A & Figure S6A) compared to the background signals from p53 CUT&RUN in *Trp53^-/-^* cells and the non-specific IgG signal in *Trp53^R172H/-^* cells. Peaks derived from two different p53^R172H^ antibodies overlapped significantly (Figure S6B), primarily occupying gene promoters (Figure S6C). Even when analyzing unique peaks identified by each antibody, the CUT&RUN signals from both correlated well (Figure S6D), suggesting that the incomplete overlap may result from the thresholds used in peak calling. Interestingly, p53^R172H^ binding was not confined to the Cxcl1 promoter; we identified even stronger peaks upstream of the Cxcl1 gene. Since these regions could function as transcriptional regulatory elements influencing Cxcl1 expression, we examined commonly used markers for such elements in *Trp53^R172H/-^* and *Trp53^-/-^* cells. We detected prominent signals for the transcription co-factor p300 and H3K27Ac (Figure 4B). Additionally, the lower levels of H3K4me3 and higher levels of H3K27Ac at these loci, compared to the Cxcl1 promoter, suggest that these regions are likely enhancers.

**Figure 4.**
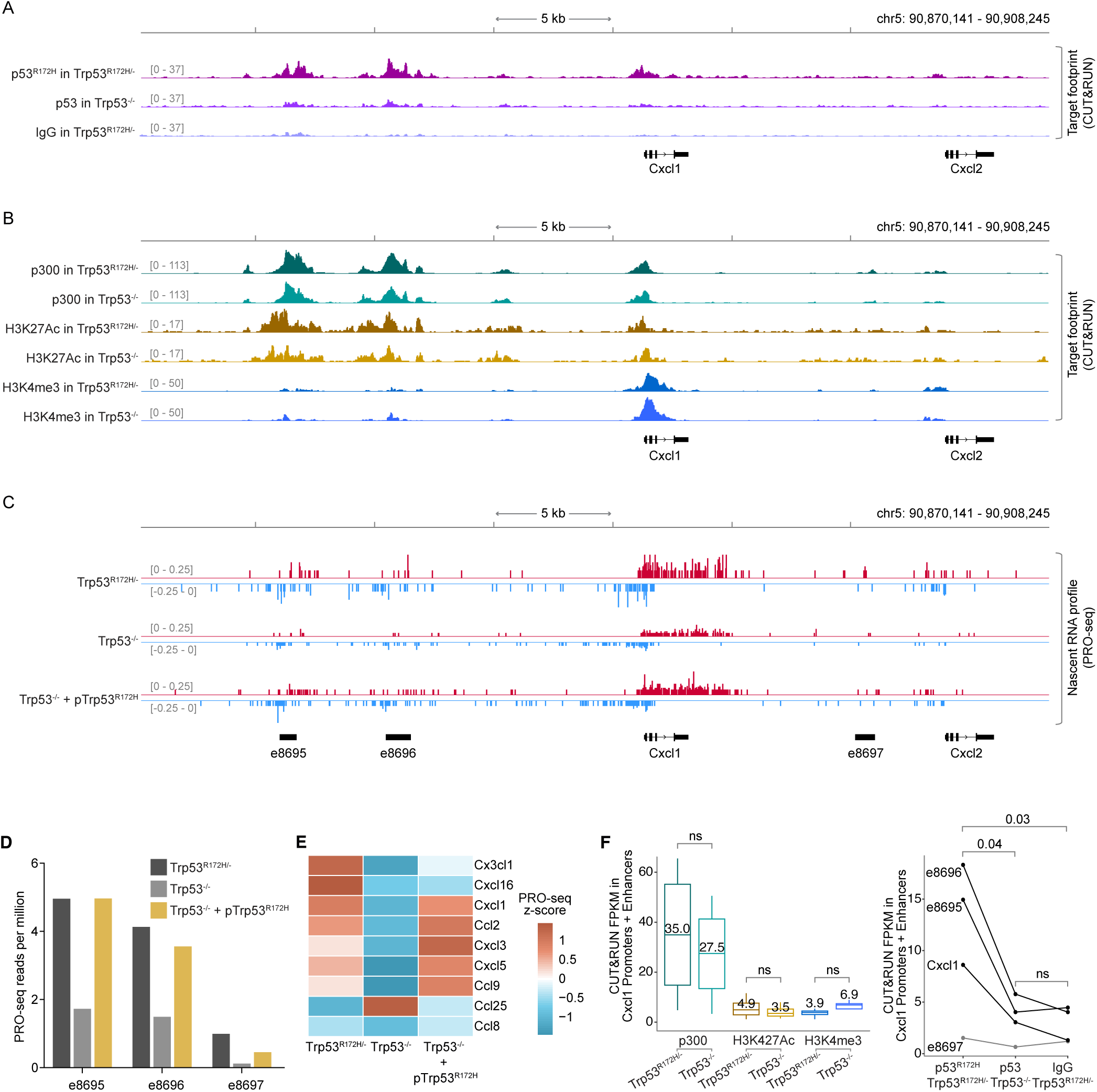
p53^R172H^ occupies and modulates Cxcl1 enhancers. **A,** p53^R172H^ occupancy at and around the *Cxcl1* gene in *Trp53^R172H/-^* and *Trp53^-/-^* cells using CUT&RUN. Non-specific IgG in *Trp53^R172H/-^*cells is used as a control. **B,** p300, H3K27Ac, and H3K4me3 occupancy at and around the *Cxcl1* gene in *Trp53^R172H/-^* and *Trp53^-/-^* cells using CUT&RUN. **C**, Nascent RNA profiles at and around the *Cxcl1* gene in *Trp53^R172H/-^*, *Trp53^-/-^*, or *Trp53^-/^*^-^ + *pTrp53^R172H^* cells using PRO-seq. Nascent RNA profiles are used to call *de novo* enhancers using dREG^71^, and the annotated enhancers are shown at the bottom (names begin with “e”). **D**, Quantification of enhancer RNA (eRNA) from PRO-seq in the three enhancers around the *Cxcl1* gene. **E**, Nascent RNA levels of the expressed Chemokine genes shown as the z-cores heatmap of PRO-seq RPM in the three isogenic cells. **F,** Quantification of p300 and histone modification levels (left) and p53^R172H^ occupancy (right) at the promoter and enhancers of the Cxcl1 gene using CUT&RUN. P-values are calculated excluding e8697 using a paired t-test.

A strong indicator of an active enhancer is the presence of short, unstable, non-polyadenylated enhancer RNA (eRNA) generated from the divergent transcription of enhancers^70^. To confirm that the p53^R172H^-occupied regions around the Cxcl1 gene are enhancers, we performed nascent RNA sequencing using PRO-seq on *Trp53^R172H/-^*, *Trp53^-/-^*, and *Trp53^-/-^* + *pTrp53^R172H^*cells (Figure 4C). We observed prominent divergent transcription at these putative enhancers. Using dREG^71^, which identifies *de novo* enhancers based on divergently transcribed nascent RNA profiles, we discovered three enhancers (e8695, e8696, and e8697, measuring 705 bp, 1065 bp, and 819 bp, respectively) within 15 kb of the Cxcl1 gene (Figure 4C). It is well-established that the level of enhancer transcription positively correlates with the transcription of target genes^70^. We quantified the eRNA at these enhancers and found that the deletion of *Trp53^R172H^* reduced their activity, while ectopic expression of p53^R172H^ mostly restored it (Figure 4D), suggesting that p53^R172H^ binding at these enhancers may significantly influence the transcription output of target genes. We measured chemokine gene transcription by quantifying nascent RNA in *Trp53^R172H/-^*, *Trp53^-/-^*, and *Trp53^-/-^*+ *pTrp53^R172H^* cells. We observed increased nascent transcription of a similar subset of chemokine genes in the presence of p53^R172H^ (Figure 4E), indicating transcriptional regulation by p53^R172H^.

To confirm the specificity of the p53^R172H^ occupancy and to ensure that the p53^R172H^ CUT&RUN signals are not a result of p53 antibodies non-specifically associating with chromatin-associated factors, we quantified p53^R172H^, p300, H3K27Ac, and H3K4me3 signals. At the promoters and enhancers of Cxcl1 (Figure 4F) and also at all p53^R172H^ CUT&RUN peaks (n=7,721) (Figure S6E & Figure S6F), we found that the enrichment of p53^R172H^ occupancy in *Trp53^R172H/-^*cells compared to *Trp53^-/-^* cells was significantly higher than the difference in the levels of chromatin-associated factors. Instead, the levels of p300 and histone modifications at the p53^R172H^- occupied regions were relatively similar between *Trp53^R172H/-^* and *Trp53^-/-^* cells. This suggests that p53^R172H^ is not responsible for opening and priming the chromatin for transcription. Instead, p53^R172H^ augments the transcription activation potential of these regions that seem to be already open and primed, likely by other transcription factors. It is plausible that p53^R172H^ commutes to these enhancers through interaction with other transcription factors. This model is consistent with a lack of a p53^R172H^-specific DNA binding motif under the p53^R172H^ CUT&RUN peaks. These observations also highlight that p300 binding and histone modifications are not the direct temporal determinants of the regulatory activity of promoters and enhancers^72^. Instead, transcription factor occupancy and divergent nascent transcription more precisely predict the activity of enhancers and the expression of their target gene^73^.

To confirm the specificity of p53^R172H^ occupancy and ensure that the p53^R172H^ CUT&RUN signals were not due to non-specific association of p53 antibodies with chromatin-associated factors, we quantified p53^R172H^, p300, H3K27Ac, and H3K4me3 signals. At both the promoters and enhancers of Cxcl1 (Figure 4F) and across all p53^R172H^ CUT&RUN peaks (n=7,721) (Figure S6E & Figure S6F), we observed that the enrichment of p53^R172H^ occupancy in *Trp53^R172H/-^* cells compared to *Trp53^-/-^* cells was significantly higher than the difference in the levels of chromatin-associated factor. The levels of p300 and histone modifications at p53^R172H^-occupied regions were relatively similar between *Trp53^R172H/-^* and *Trp53^-/-^*cells, suggesting that p53^R172H^ is not responsible for opening and priming the chromatin for transcription. Instead, p53^R172H^ appears to enhance the transcriptional activation potential of regions that are likely already open and primed by other transcription factors. It is plausible that p53^R172H^ is recruited to these enhancers through interactions with other transcription factors. These observations also suggest that p300 binding and histone modifications are not direct temporal determinants of the regulatory activity of promoters and enhancers^72^. Instead, transcription factor occupancy and divergent nascent transcription more accurately predict enhancer activity and the expression of their target genes^73^.

### Cxcl1 enhancers dictate Cxcl1 expression and its immunosuppressive function

To test whether and to what extent the Cxcl1 enhancers regulate the Cxcl1 expression, we systematically deleted them individually or in pairs using CRISPR-Cas9 (Figure 5A). We observed that the deletion of e8695 and e8696 but not e8697 (denoted as Δe8695, Δe8696, and Δe8697, respectively) significantly reduced the Cxcl1 expression (Figure 5B). These enhancer-deleted cells are *Kras^G12D/+^*, *Trp53^R172H/-^*, and *Cxcl1^+/+^*. The dual deletion of e8695 and e8696 (Δe8695 + Δe8696) further reduced the Cxcl1 expression, but the additional reduction was modest. To examine their contribution to the PDAC growth and TME, we orthotopically implanted the Δe8695, Δe8696, and Δe8697 cells in the pancreas of immunocompetent mice and extracted the tumors after 21 days. Δe8695 and Δe8696 tumors were significantly and consistently reduced in size as compared to their parental cell, and the extent of their effect was similar to their impact on Cxcl1 levels, whereas Δe8697 tumor size was partially reduced in one cohort but not detectably reduced in another cohort (Figure 5C & Figure S7A).

**Figure 5.**
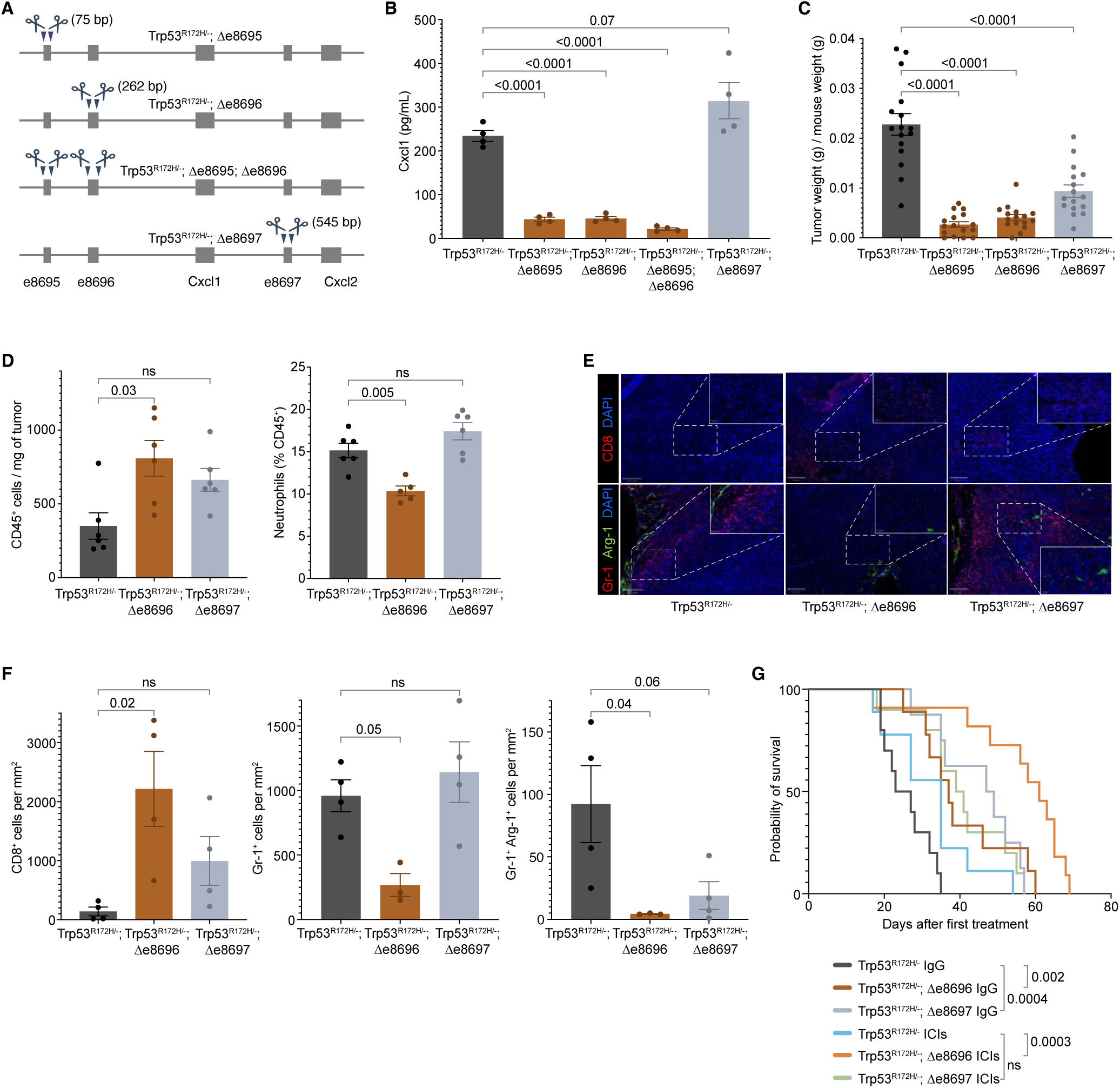
p53^R172H^-occupied Enhancers regulate Cxcl1 expression and dictate Cxcl1- mediated immunosuppression. **A,** Generation of enhancer-deleted isogenic cells from the parental *Trp53^R172H/-^*cells. Two enhancers upstream of the *Cxcl1* gene and one downstream of the *Cxcl1* gene are deleted individually or in a pair using CRISPR/Cas9. The size of deleted regions is indicated in parentheses. **B,** Quantification of the Cxcl1 chemokine level in enhancer-deleted isogenic cells and comparison with the *Trp53^R172H/-^* cells. P-values are calculated from a one-way ANOVA test followed by a post hoc test with Benjamini-Hochberg correction. **C,** Weights of the enhancer-deleted isogenic tumors compared with the *Trp53^R172H/-^*tumors. *Trp53^R172H/-^*;Δe8695;Δe8696 cells were not implanted in mice due to the minimal effect over either *Trp53^R172H/-^*;Δe8695 or *Trp53^R172H/-^*;Δe8696 cells. P-values are calculated from a one-way ANOVA test followed by a post hoc test with Benjamini-Hochberg correction. **D,** Effect of the e8696 and e8697 status in leukocyte and neutrophil infiltration in PDAC tumors. P-values are calculated from a one-way ANOVA test followed by a post hoc test with Benjamini-Hochberg correction. **E,** Representative multiplex immunofluorescence staining of *Trp53^R172H/-^*;Δe8695 and *Trp53^R172H/-^*;Δe8696 tumors. The scale bars correspond to 200 μm and 50 μm in the main view and the magnified view, respectively. **F,** Quantification of immune cell density from randomly selected 500 x 500 µm regions. P-values are calculated from a one-way ANOVA test followed by a post hoc test with Benjamini-Hochberg correction. **G,** Kaplan-Meier survival curves showing the effects of e8696 or e8697 status and ICIs in the survival of mice implanted with parental *Trp53^R172H/-^* or *Trp53^R172H/-^;*Δe8696, or *Trp53^R172H/-^;*Δe8697 cells. Control mice were treated with IgG. *Trp53^R172H/-^* mice survival cohort is the same as in Figure S4D & Figure 3I. P-values are calculated using a log-rank (Mantel-Cox) test.

As shown in Figure 3, the ablation of Cxcl1 alone reprogrammed the immune landscape of the PDAC TME. Although Δe8695 and Δe8696 reduced Cxcl1 expression and tumor size, we sought to ensure that enhancer deletion recapitulated the Cxcl1-mediated immune reprogramming. To minimize batch effects, we processed all mice and samples together, which limited us to two enhancer-deleted cell lines. We selected Δe8696 and Δe8697 due to their equidistant positions from the Cxcl1 gene and their variability in inducing Cxcl1 expression. Moreover, based on Cxcl1 expression and tumor weight (Figure 5B & Figure 5C), we anticipated that Δe8695 would recapitulate the results of Δe8696. Consequently, we examined the immune landscape, tumor growth, and survival with Δe8696 and Δe8697 cells. As seen in the *Cxcl1^-/-^* tumors, we observed higher infiltration of immune cells, a lower fraction of neutrophils, and higher innate lymphoid cells, macrophages, and NK cells in Δe8696 tumors (Figure 5D & Figure S7B). We used immunofluorescence to further examine the immune composition of the tumors (Figure 5E). We found a higher density of cytotoxic T cells (CD8^+^) and a lower density of immunosuppressive neutrophils (Gr-1^+^) and MDSCs (Gr-1^+^Arg-1^+^) in Δe8696 tumors compared to *Trp53^R172H/-^* tumors (Figure 5F).

We then tested if the reprogrammed immune microenvironment influenced the efficacy of ICIs. We observed the mice with Δe8696 tumors responded well to ICIs (Figure 5G). The survival and ICIs benefits were not observed in Δe8697 despite its similar proximity to the Cxcl1 gene. e8697 is closer to the Cxcl2 gene, which is lowly expressed in these cells. Moreover, the transcription of e8697 is lower than that of e8696, and the CUT&RUN signal of p53^R172H^ is not detected at this enhancer (Figure 4A & Figure S6A). These observations suggest that the p53^R172H^-occupied e8696 is the critical regulator of Cxcl1 expression and p53^R172H^-mediated immunosuppression in PDAC.

### NF-κB occupies Cxcl1 promoter and enhancers

The arginine residue at position 172 of p53 stabilizes the DNA binding domain by contacting a zinc ion^18^. Therefore, the common R172H mutation alters p53’s conformation and abolishes its canonical DNA binding activity^17^. To further understand the mechanism underlying p53^R172H^ DNA association, we investigated whether p53^R172H^ associates with DNA independently or relies on other transcription factors. Our analysis of the DNA sequences under the p53^R172H^ CUT&RUN peaks revealed no enrichment of the canonical wild-type p53 DNA binding motif or any novel motif. Instead, we observed an enrichment of several known transcription factors under the p53^R172H^ peaks.

When we specifically examined the enhancers and promoters of the p53^R172H^-dependent chemokines, we found the NF-κB motif to be the most enriched, present in approximately 70% of examined sites (Figure 6A). To test if NF-κB binds these putative NF-κB sites, we used CUT&RUN with an antibody against the RelA (p65) subunit of NF-κB. We observed prominent NF-κB occupancy at the Cxcl1 promoter and enhancers (Figure 6B). Quantification showed a ∼20-25% decrease in NF-κB occupancy in *Trp53^-/-^* cells compared to *Trp53^R172H/-^*cells (Figure 6C), suggesting a role of p53^R172H^ in increasing NF-κB occupancy at these sites. Despite similar p300 occupancy and histone modifications (Figure S8A), the average NF-κB CUT&RUN signal was similarly lower in *Trp53^-/-^* cells compared to *Trp53^R172H/-^* cells (Figure 6D).

**Figure 6.**
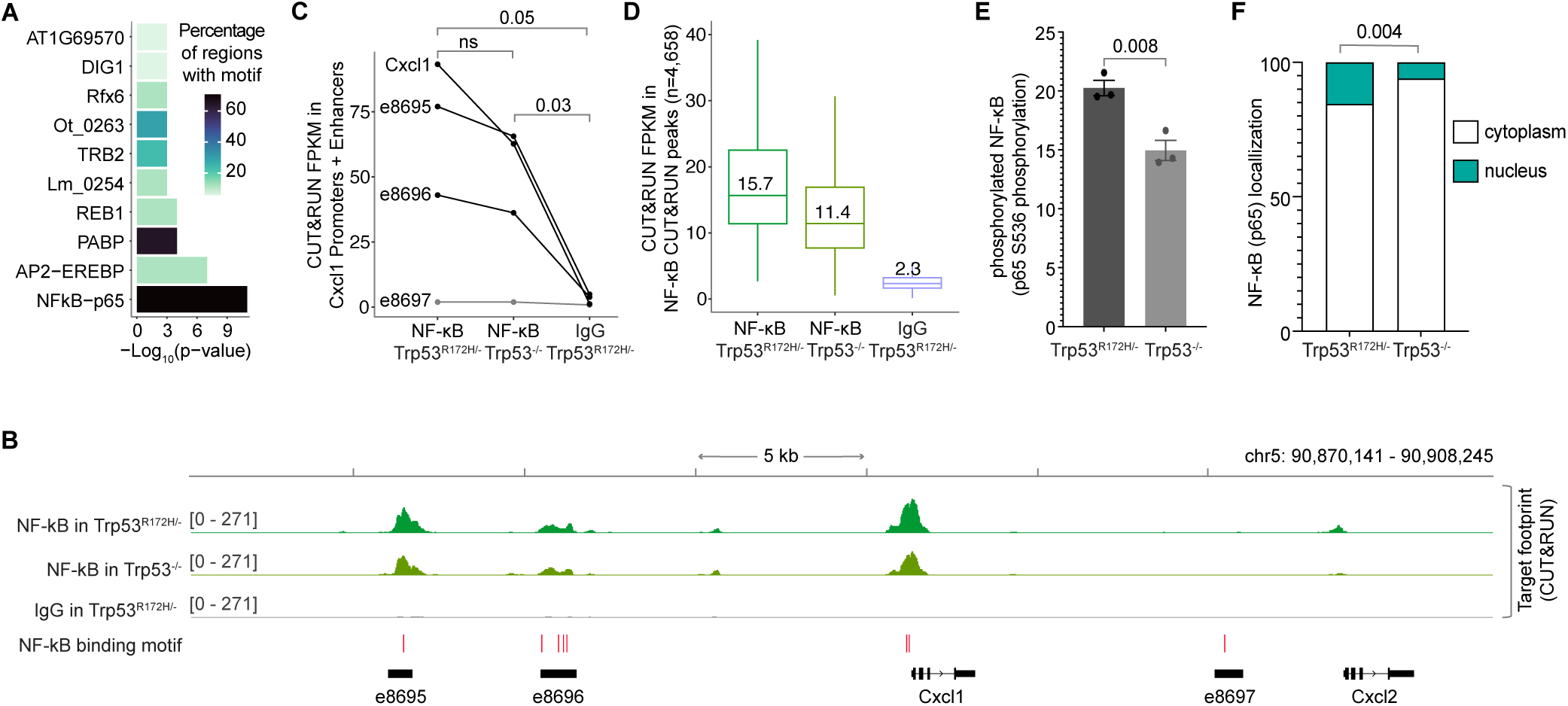
p53^R172H^ occupancy in Cxcl1 Enhancers is dependent on NF-κB. **A,** Enriched transcription factor motifs at the promoters and enhancers of the p53^R172H^- dependent chemokine genes identified using HOMER^105^. **B**, NF-κB occupancy at the *Cxcl1* gene and enhancers in *Trp53^R172H/-^* and *Trp53^-/-^* cells using CUT&RUN. Non-specific IgG is used as a control. The HOMER-identified NF-κB motifs are shown at the bottom (red bars). **C**, Quantification of NF-κB occupancy at the promoter and enhancers of the Cxcl1 gene using CUT&RUN. P-values are calculated excluding e8697 using a paired t-test. **D**, Quantification of NF-κB occupancy at the NF-κB CUT&RUN peaks using CUT&RUN. **E**, Quantification of NF-κB phosphorylation in the *Trp53^R172H/-^* and *Trp53^-/-^* cells using ELISA on cell extracts. The antibody targets S536 in the p65 subunit of NF-κB. **F**, Quantification of NF-κB localization in the cytoplasm vs nucleus using western blot (serial dilution of protein lysates). Vinculin and histone H3 were used as the markers of cytoplasmic and nuclear fractions, respectively. P-values are calculated from a two-way ANOVA test.

We observed a similar trend of lower NF-κB occupancy in *Trp53^-/-^* cells compared to *Trp53^R172H/-^* cells in the p53^R172H^ CUT&RUN peaks (Figure S8B). This lower occupancy was not due to differential expression of NF-κB between the two cell types (Figure S8C). Instead, we found that the phosphorylation of the RelA subunit of NF-κB and its nuclear localization is reduced by ∼25% in the absence of p53^R172H^ (Figure 6E & Figure 6F). Phosphorylated NF-κB translocates into the nucleus, where it binds and regulates the expression of pro-inflammatory genes^74^. These data suggest that while NF-κB expression is unaffected by p53^R172H^, its phosphorylation, subsequent nuclear localization, and genome occupancy are moderately increased in the presence of p53^R172H^. However, this ∼20-25% increase in NF-κB occupancy at the promoters and enhancers of Cxcl1 does not fully explain the four-fold higher expression of Cxcl1 in *Trp53^R172H/-^* cells compared to *Trp53^-/-^* cells (Figure 1F).

Our observation of both p53^R172H^ and NF-κB occupancy at the Cxcl1 enhancers and promoters led us to examine the overlap between p53^R172H^ and NF-κB CUT&RUN peaks. We found a significant overlap in their genomic occupancy, with most co-occupied regions coinciding with known promoters and enhancers (Figure S8D). We quantified p53^R172H^ and NF-κB occupancy, p300 levels, histone modifications, and nascent transcription across three sets of CUT&RUN peaks: unique p53^R172H^ peaks, common peaks, and unique NF-κB peaks (Figure S9A-C). In each case, the levels of chromatin-associated factor were similar between *Trp53^R172H/-^* and *Trp53^-/-^* cells. However, differences in p53^R172H^ occupancy were reflected in nascent RNA profiles, indicating that nascent transcription is a better indicator of the transcriptional state of promoters and enhancers than chromatin-associated factors. These findings demonstrate that NF-κB binds to the promoters and enhancers of Cxcl1 largely independently of p53^R172H^, and its genome-wide occupancy significantly overlaps with that of p53^R172H^.

### p53^R172H^ depends on NF-κB for Cxcl1 enhancer binding

We next investigated whether p53^R172H^ genome occupancy depends on NF-κB. To test this, we examined the impact of TPCA-1, a potent IKKβ inhibitor that prevents NF-κB activation, on NF- κB and p53^R172H^ occupancy using CUT&RUN. As expected, TPCA-1 treatment significantly reduced NF-κB binding at the Cxcl1 enhancers in both *Trp53^R172H/-^* and *Trp53^-/-^* cells (Figure 7A). More importantly, p53^R172H^ occupancy at the Cxcl1 enhancers was similarly reduced following TPCA-1 treatment, strongly suggesting that p53^R172H^ binding is dependent on the NF-κB pathway.

**Figure 7.**
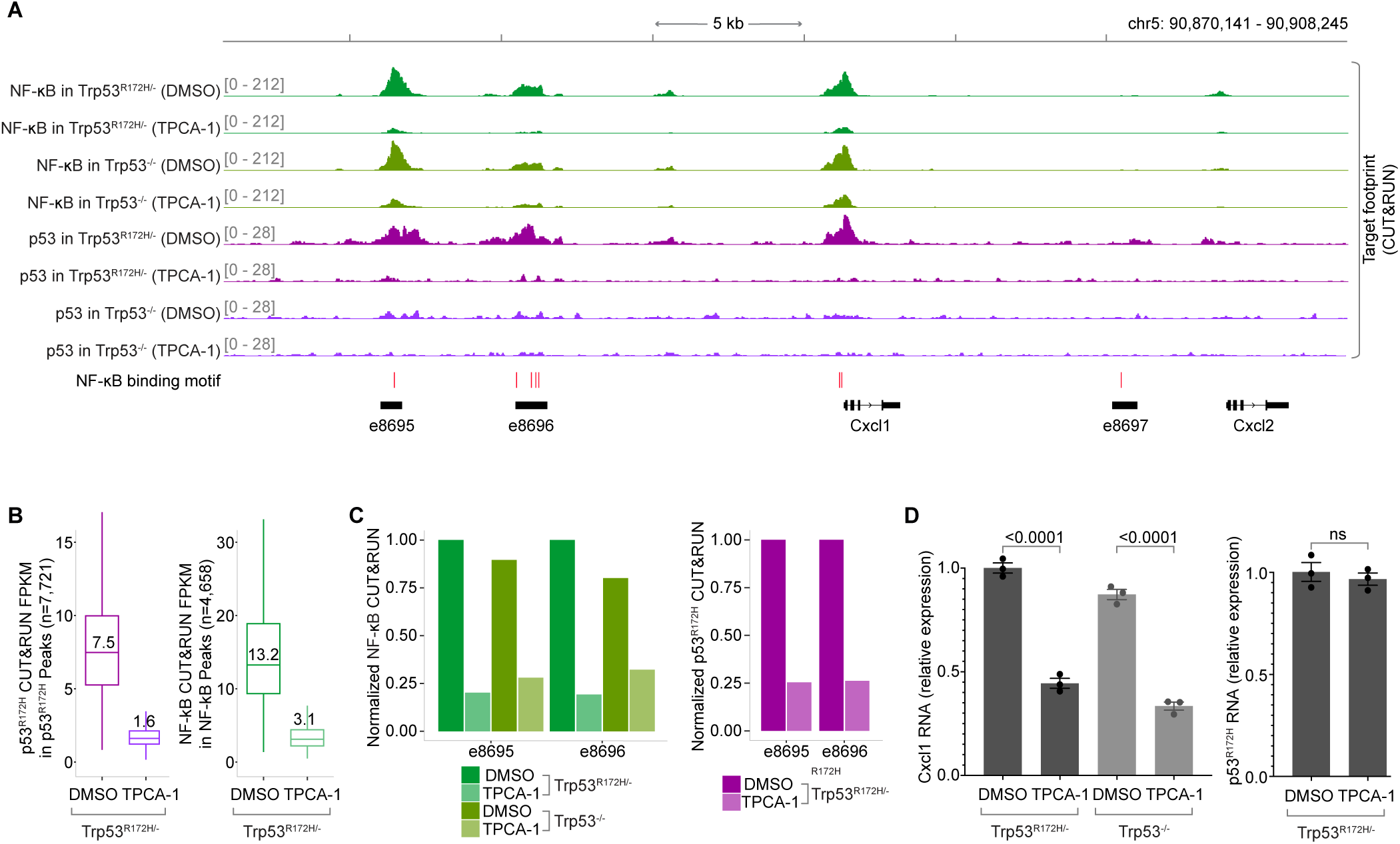
NF-κB inhibition abrogates p53^R172H^ occupancy. **A**, Effect of inhibiting NF-κB activation (5 uM TPCA-1) in NF-κB and p53^R172H^ occupancy at the *Cxcl1* gene and enhancers. DMSO treatment serves as a control. **B**, Change in p53^R172H^ occupancy (left) and NF-κB occupancy (right) by NF-κB inhibition in the p53^R172H^ and NF-κB CUT&RUN peaks, respectively. **C**. Effects of TPCA-1 treatment in NF-κB (left) and p53^R172H^ (right) occupancy in the two detectable enhancers of the *Cxcl1* gene. **D**, Effects of TPCA-1 treatment in *Trp53^R172H^* (left) and *Cxcl1* (right) mRNA levels using RT- qPCR.

Our data showed that CUT&RUN signals for all p53^R172H^ peaks and nearly all NF-κB peaks (except one) decreased following NF-κB inhibition by TPCA-1 (Figure 7B & Figure S10A), highlighting the broad impact of NF-κB on p53^R172H^ chromatin binding. Specifically, the TPCA-1 treatment reduced the occupancy of both NF-κB and p53^R172H^ to approximately 20% at the Cxcl1 enhancers, illustrating the critical role of NF-κB in facilitating p53^R172H^ recruitment to these sites (Figure 7C). Additionally, we observed that TPCA-1 treatment led to a decrease in Cxcl1 gene expression, as expected, but did not affect *Trp53^R172H^* gene expression. This finding suggests that the reduced occupancy of p53^R172H^ was not due to lower *Trp53^R172H^* expression but rather the direct effect of NF-κB pathway inhibition (Figure 7D).

Collectively, our findings reveal a novel mechanism by which the cancer-cell-intrinsic p53^R172H^ occupies the enhancers of the Cxcl1 gene in an NF-κB-dependent manner. This appropriation of the NF-κB pathway by p53^R172H^ amplifies Cxcl1 expression, contributing to the establishment of an immunosuppressive TME that, in turn, diminishes the efficacy of ICIs. These insights highlight potential therapeutic targets for enhancing ICI efficacy in PDACs harboring *Trp53* missense mutations and elevated Cxcl1 expression.

## Discussion

Our findings indicate that p53^R172H^ promotes tumor growth by modulating cancer-cell-specific gene expression programs that shape the TME to suppress anti-tumor immunity. Specifically, *Trp53^R172H/-^*tumors exhibit fewer T cells, higher MDSC infiltration, and reduced ICI efficacy in mouse models of PDAC. At the molecular level, p53^R172H^ exerts its immunosuppressive effects primarily through the regulation of chemokine genes, particularly Cxcl1. p53^R172H^ binds to distal enhancers of Cxcl1 and amplifies its expression in an NF-κB-dependent manner. This interaction suggests a broader mechanism by which mutant p53 may co-opt enhancers—vastly outnumbering gene promoters—by interacting with the enhancer’s cognate binders to modulate cell-intrinsic gene expression, ultimately influencing cell-extrinsic features. This study not only elucidates the specific mechanisms of immunosuppression facilitated by p53^R172H^ but also suggests broader implications for p53 missense mutations and their cooperation with other transcription factors in cancer progression.

Enhancers have emerged as key players in disease pathology. Over 90% of disease-associated variants are located in non-coding regions, predominantly within enhancers^75^. Despite this recognition, understanding the precise mechanisms by which enhancers influence disease remains a challenge. However, recent advances, exemplified by the FDA approval of Casgevy, the first CRISPR-Cas9-based therapy targeting the enhancer regulating the BCL11A transcription factor for sickle cell anemia^76^, underscore the therapeutic potential of manipulating enhancer activity. Our study further illuminates the significance of enhancers, revealing how oncogenic factors can appropriate them to drive the expression of immunosuppressive chemokines for PDAC progression and confer resistance to ICIs.

Our findings hold two other significant implications. First, they offer mechanistic insights into the therapeutic targeting of immunosuppressive chemokines and their receptors, as well as the upstream signaling pathways, in cancer treatment. High levels of Cxcl1 are associated with poor prognosis in various cancers, including PDAC^77^. A similar dependence of selected chemokine genes on p53^R172H^ was observed in previous studies. shRNA-mediated knockdown of mutant p53 in human colon adenocarcinoma cell line SW480 (harboring R273H/P309S mutations) and pancreatic cancer cell line MIA-PaCa-2 (harboring R248W mutation) also reduced Cxcl1 expression^78^. This supports the promise of current clinical trials testing the blockade of tumor cell-derived Cxcl1^79^, Cxcr2 deletion^80^, or its inhibition^81^. Similarly, the NF-κB signaling pathway that dictates Cxcl1 expression is activated in many cancers. However, despite its potential, no effective NF-κB inhibitor is available for clinical use in cancer treatment due to significant toxicities associated with blocking this pathway^82,83^. Our characterization of an enhancer targeting a key effector of NF-κB-mediated immunosuppression suggests a promising avenue for reducing toxicity while maintaining therapeutic efficacy.

Secondly, our findings establish a mechanistic link between a common mutation in the most frequently mutated gene and the ineffectiveness of ICIs in certain cancers. Mutations in transcription factors drive many cancers^51^, and enhancers vastly outnumber genes. The exploitation of enhancers by mutated transcription factors in our study highlights the importance of this mechanism and underscores the need for further investigation into tissue- and mutation-specific enhancer-gene regulatory circuits. It is conceivable that mutated transcription factors interact with abundant enhancers in the genome through liquid-liquid phase condensates, driving aberrant gene programs^84^.

More importantly, our observations align with clinical data. A mutant p53 RNA expression signature shows a significant correlation with reduced survival in 11 cancer types^85,86^. Similarly, PDAC with p53 missense mutations is associated with poor survival^40^, and compared to p53- null, they have enhanced fibrosis and lower lymphocyte and CD8^+^ T cell infiltration^86,87^. Degradation of mutant p53 and restoration of wild-type function are targets of several clinical trials^25,88^. Evidence suggests that the mutant p53-specific immune reprogramming in cancer is mediated by Cxcl1 overexpression. For example, ectopic expression of p53^R175H^, but not wild-type p53, whose level and activity are tightly suppressed in healthy cells^89^, increases Cxcl1 expression in colon cancer^90^. Similarly, overexpression of p53^R175H^, but not wild-type p53, specifically in the context of KRAS^G12D^ mutation, enhanced Cxcl1 expression in human pancreatic epithelial cells^30^. Another study switching p53^WT^ expression to p53^R172H^ revealed enrichment of cytokine-cytokine receptor genes, emphasizing the role of mutant p53 in cytokine regulation^43^. Moreover, in pancreatic ductal epithelial cells with Kras^G12D^ mutation, chemokine genes Cxcl2 and Cxcl5 exhibited significantly higher expression in p53^R172H^ cells compared to p53^WT^ cells^56^. It is plausible that the p53^R172H^ mutation interacts with NF-κB, utilizing it as a vehicle to reach specific enhancers. With a functional and potent transactivation domain, p53^R172H^ could elevate the expression of the Cxcl1 gene beyond the baseline induction by NF- κB alone. Prior studies have indicated mutant p53’s propensity to prolong NF-κB signaling in cancer contexts^91^, while reciprocal inhibition between p53^WT^ and NF-κB has also been documented^92,93^. Moreover, physical interaction between p53^R172H^ and NF-κB has been reported^19,20^, possibly explaining the presence of p53^R172H^ at Cxcl1 enhancers.

Our study suggests that a small fraction of PDAC patients who are p53^NULL^ may respond to ICIs. Nonetheless, combination immuno-oncology therapy in pancreatic cancer yielded only a 3.1% response rate^5^. While it is a small fraction, the genotype of PDAC patients that respond to ICIs warrants attention. It’s plausible that tumors relying on p53^R172H^-mediated immunosuppression may become vulnerable to ICIs upon abrogation of this dependence. However, we lack evidence to suggest that p53^R172H^ can serve as a biomarker for ICI responsiveness in PDAC at this stage. Instead, our data, along with evidence from the literature, suggest that overexpression of Cxcl1, implicated in various cancers through different mechanisms^94–100^, could be a better biomarker.

Our work presents a novel and potent approach to abrogate Cxcl1 expression in PDAC by targeting enhancers, which generally have higher specificity and lower toxicity.

## Supporting information

Table_1_Primer_Oligos

## ACKNOWLEDGMENTS

This research was funded by Program Project Grant P01-CA042063 from the NCI (P.A.S.), the United States Public Health Service grants R01-GM034277 from the NIH (P.A.S.), R35- CA197743, U01-CA224348, R01-CA259253, R01-CA208205, R01-NS118929, U01-CA261842, and grants from the Ludwig Cancer Center at Harvard, Nile Albright Research Foundation, Jane’s Trust Foundation, National Foundation for Cancer Research to (R.K.J.), NCI R00CA204595-05 and Pew-Stewart Scholarship to (S.S.), and Howard Hughes Medical Institute to (H.M.S.). The Emerald Foundation Postdoctoral Transition Award currently supports D.B.M. The Gertrude B. Elion Research Fellowship from GSK and the Ludwig Cancer Institute at MIT previously supported him. We thank our colleagues Tyler Jacks, William Freed-Pastor, the Late Brandon Horton, Shayla Nguyen, Skyler Kauffman, and Nathan Han for their helpful discussion. We thank the Swanson Biotechnology Center Flow Cytometry Facility for help with FACS and Stuart Levine and the staff at BioMicro Center at the Koch Institute for Integrative Cancer Research for their help with NGS. We also thank the Animal Facility at MGH and the FACS core at the Ragon Institute.

## AUTHOR CONTRIBUTIONS

Conceptualization, D.B.M., R.K.J., and P.A.S.; data acquisition, D.B.M., H.K., S.A.C., E.M., K.N., R.M., W.W.H., I.C., B.S., L.K.Y., J.F., S.K.W., V.P.C., E.M., and Y.C. ; data analysis, D.B.M., H.K., S.A.C., E.M., K.N., W.W.H., B.S., L.K.Y., A.S., and V.P.C.; funding acquisition, D.B.M., H.M.S., S.S., R.K.J., and P.A.S.; writing– original draft, D.B.M and P.A.S; writing– review and editing, all.

## KEY RESOURCES TABLE

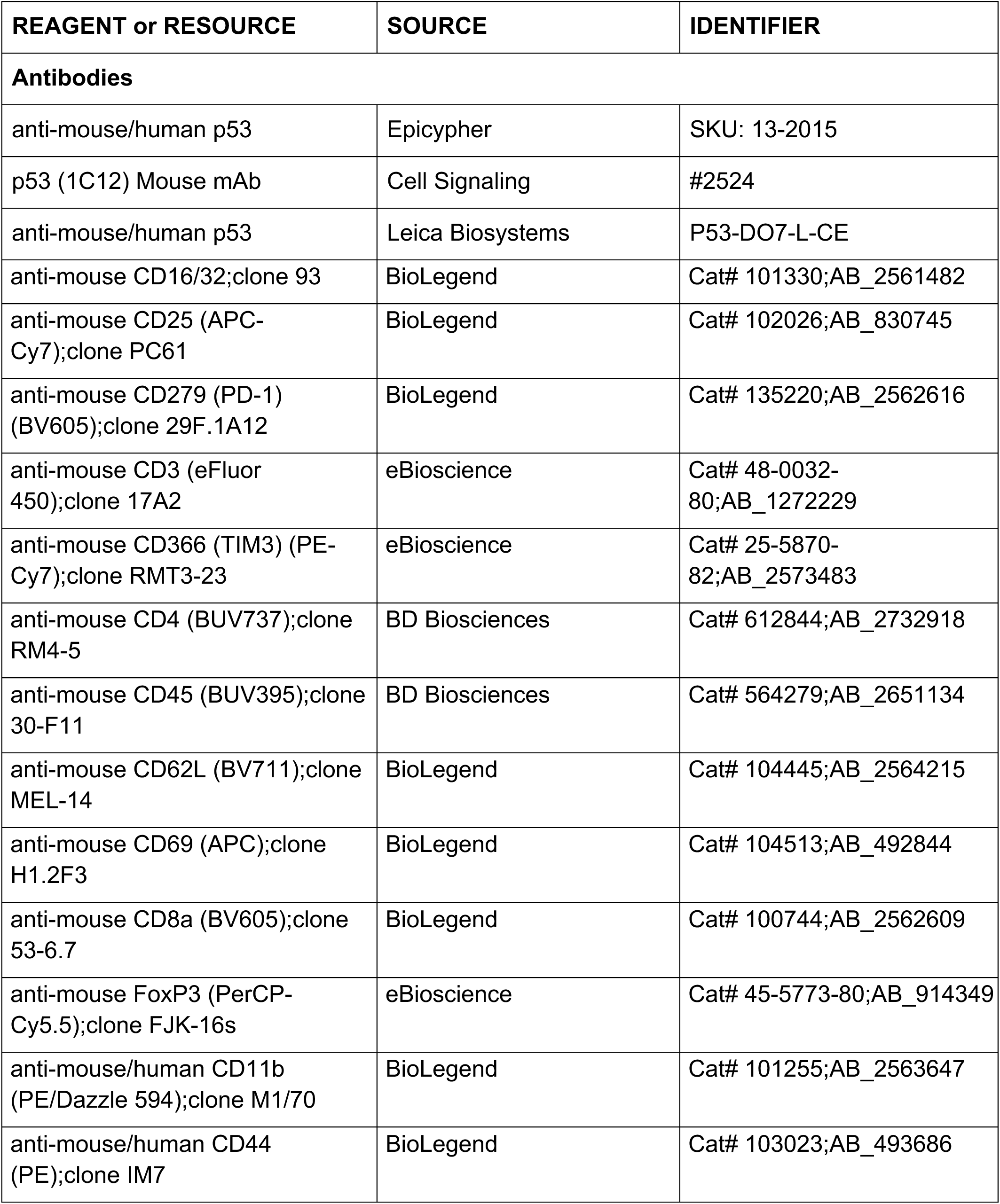
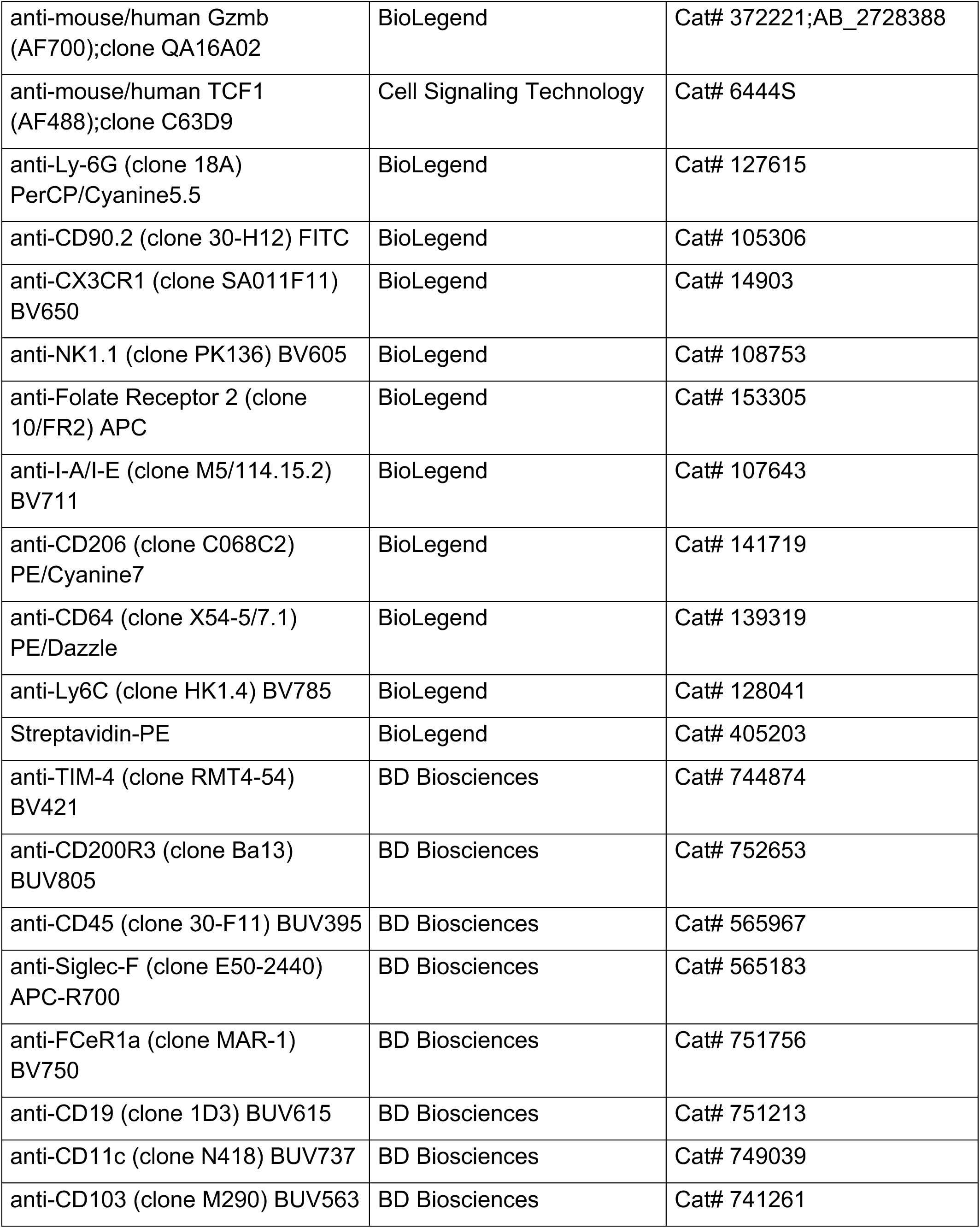
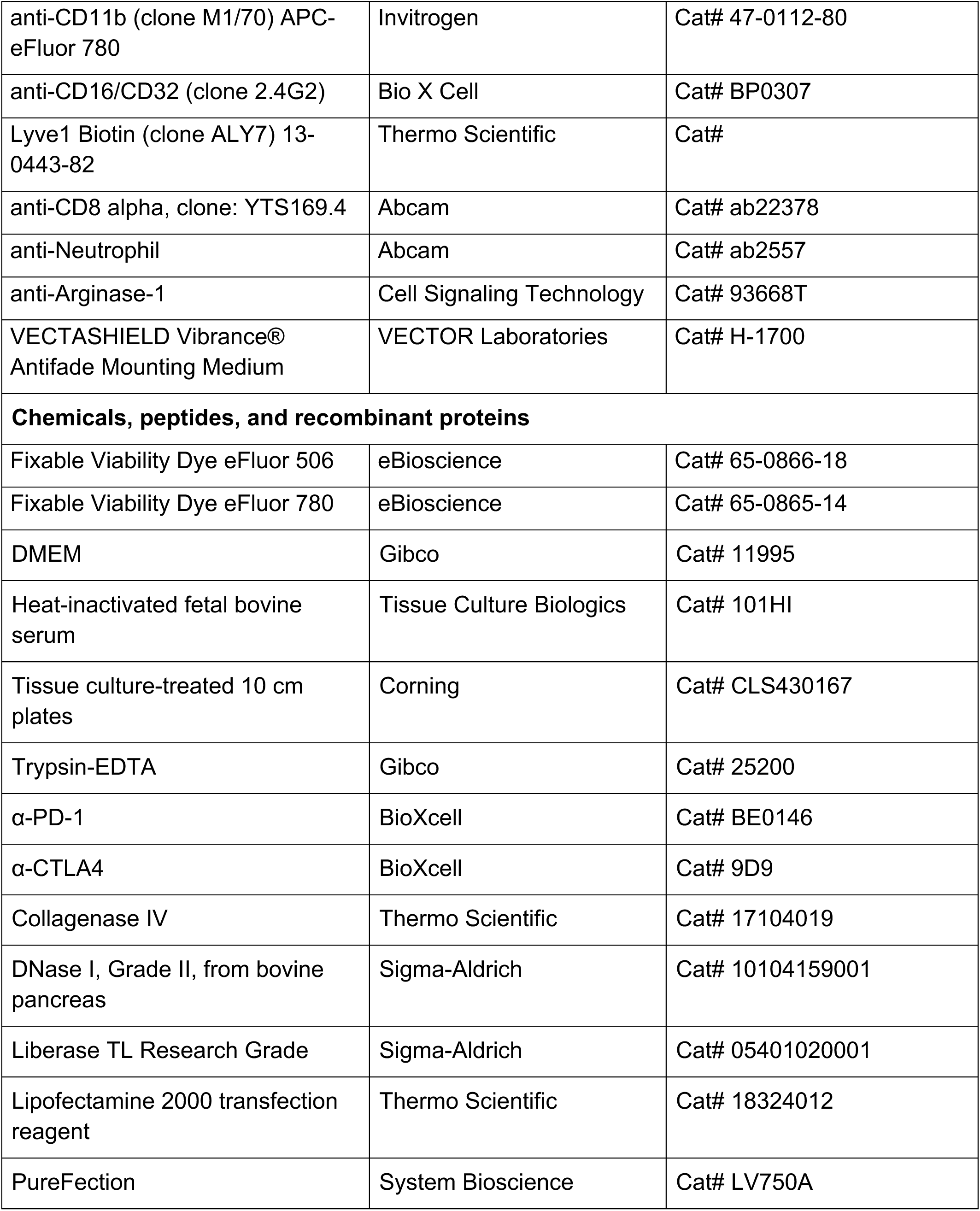
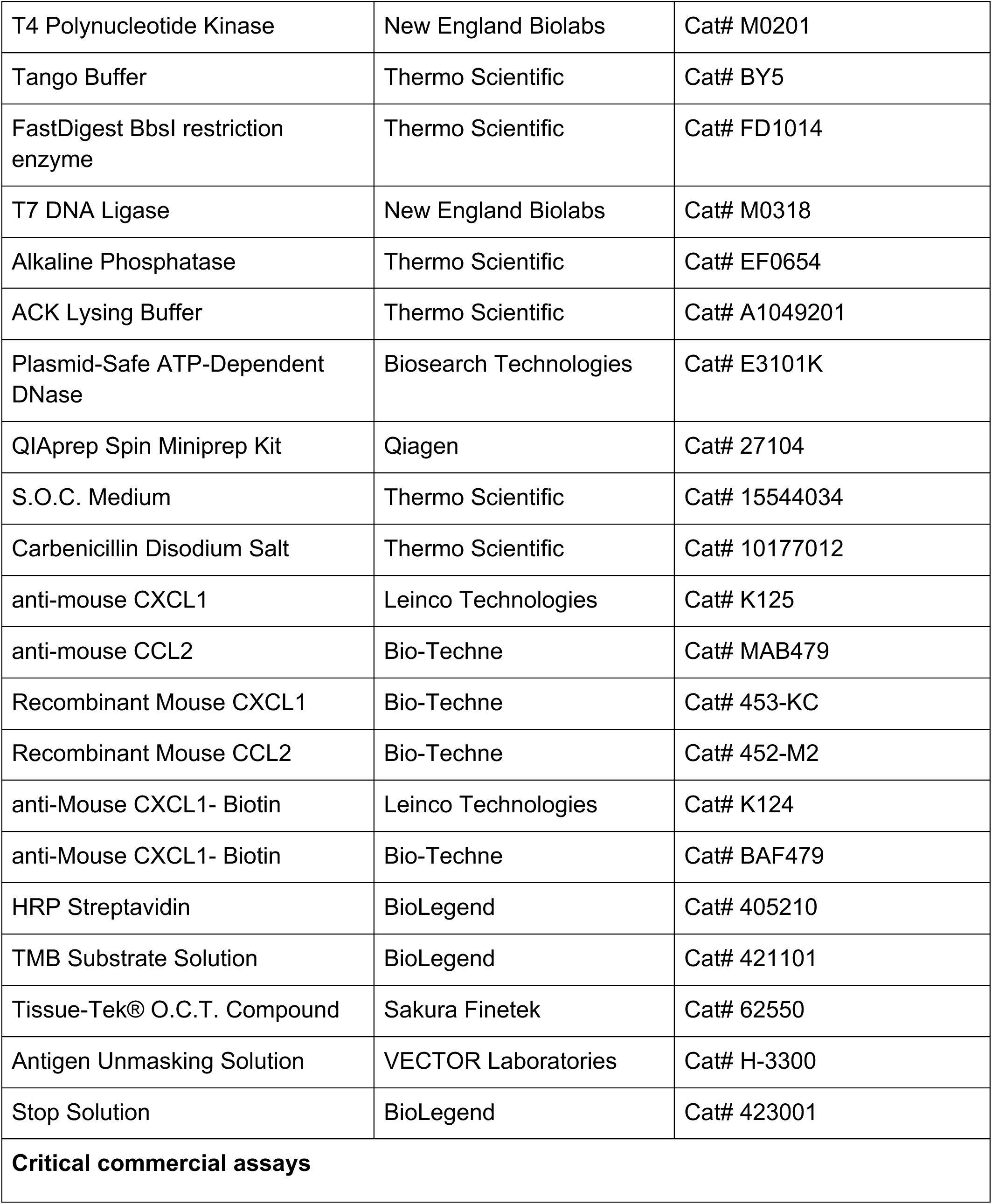
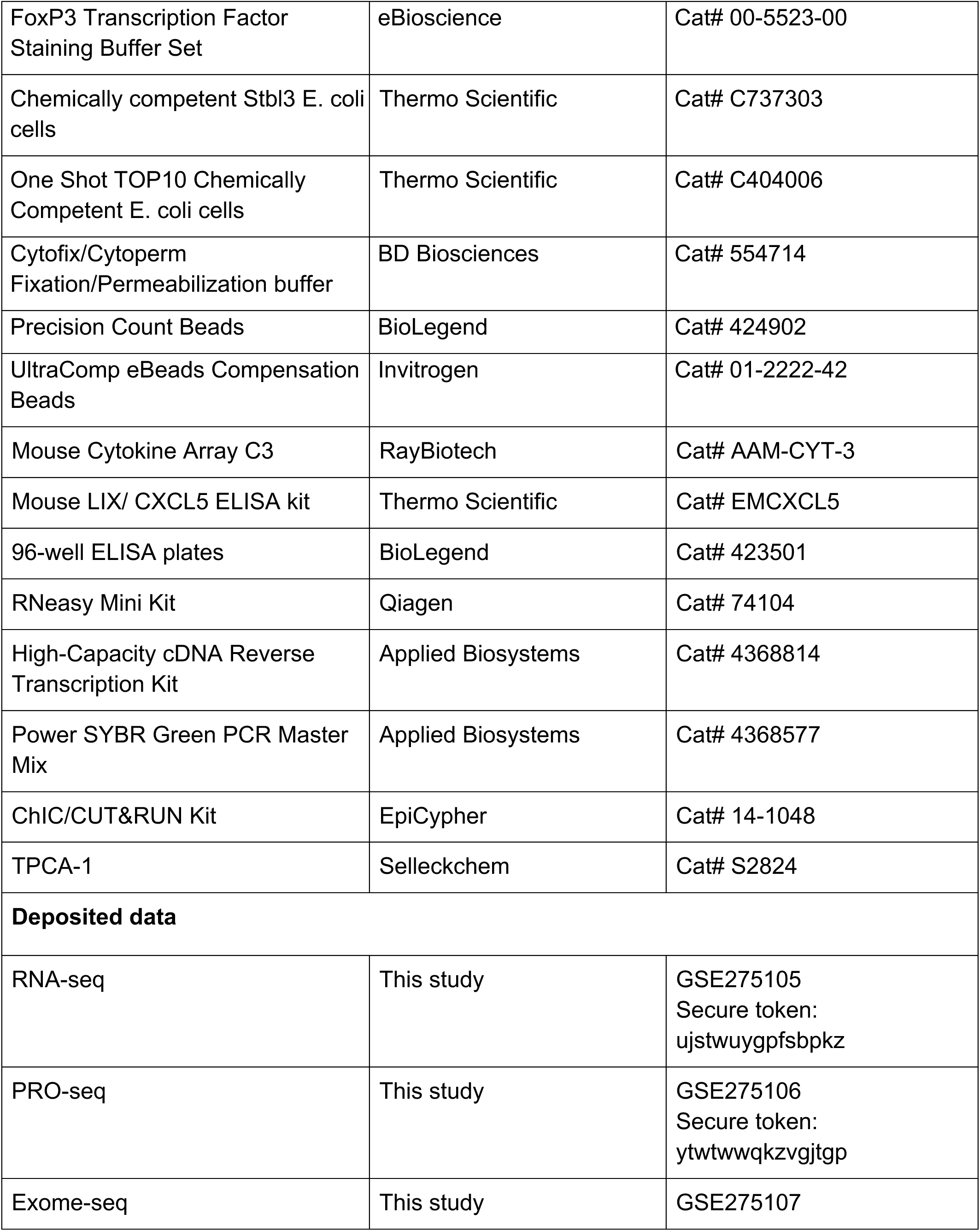
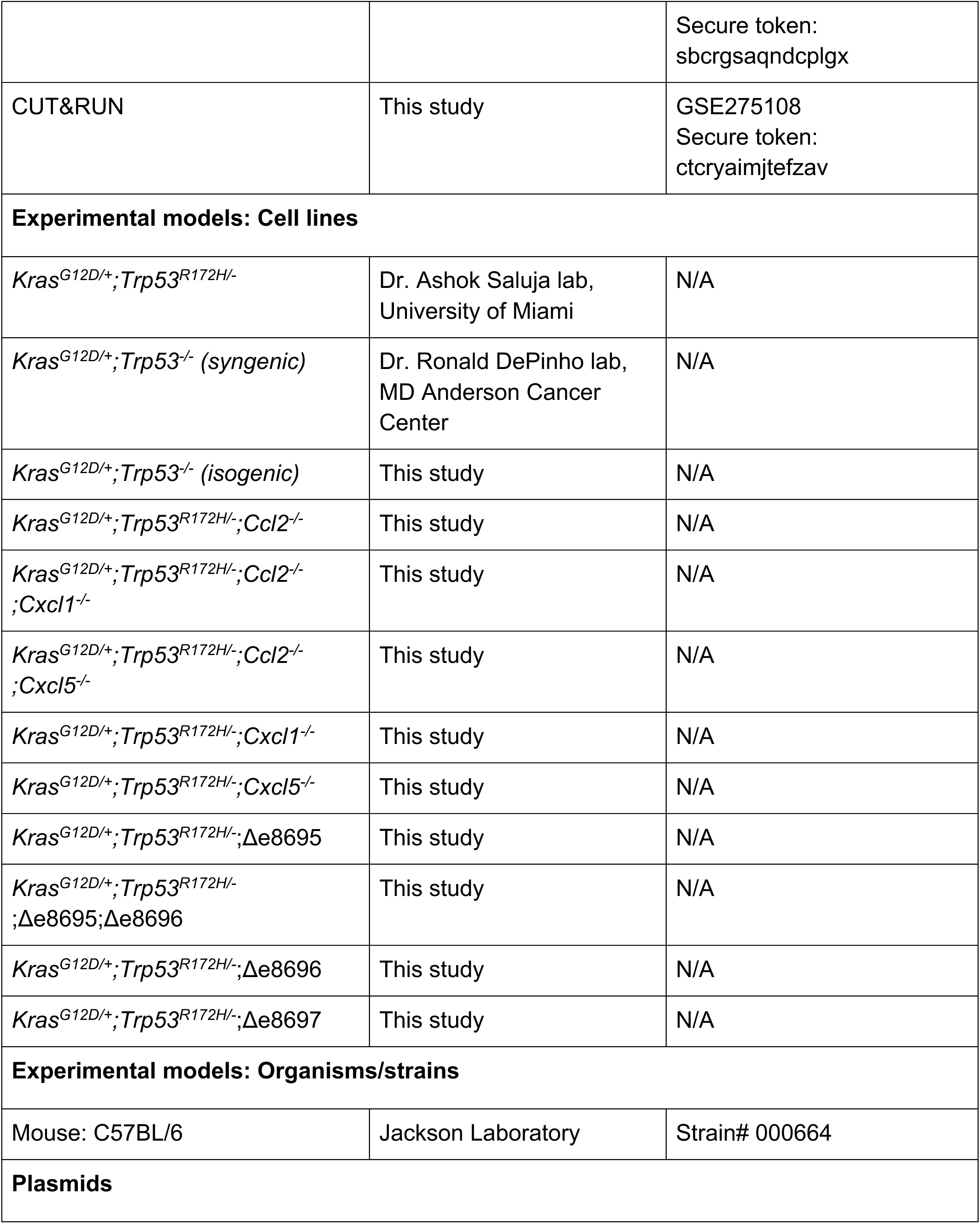
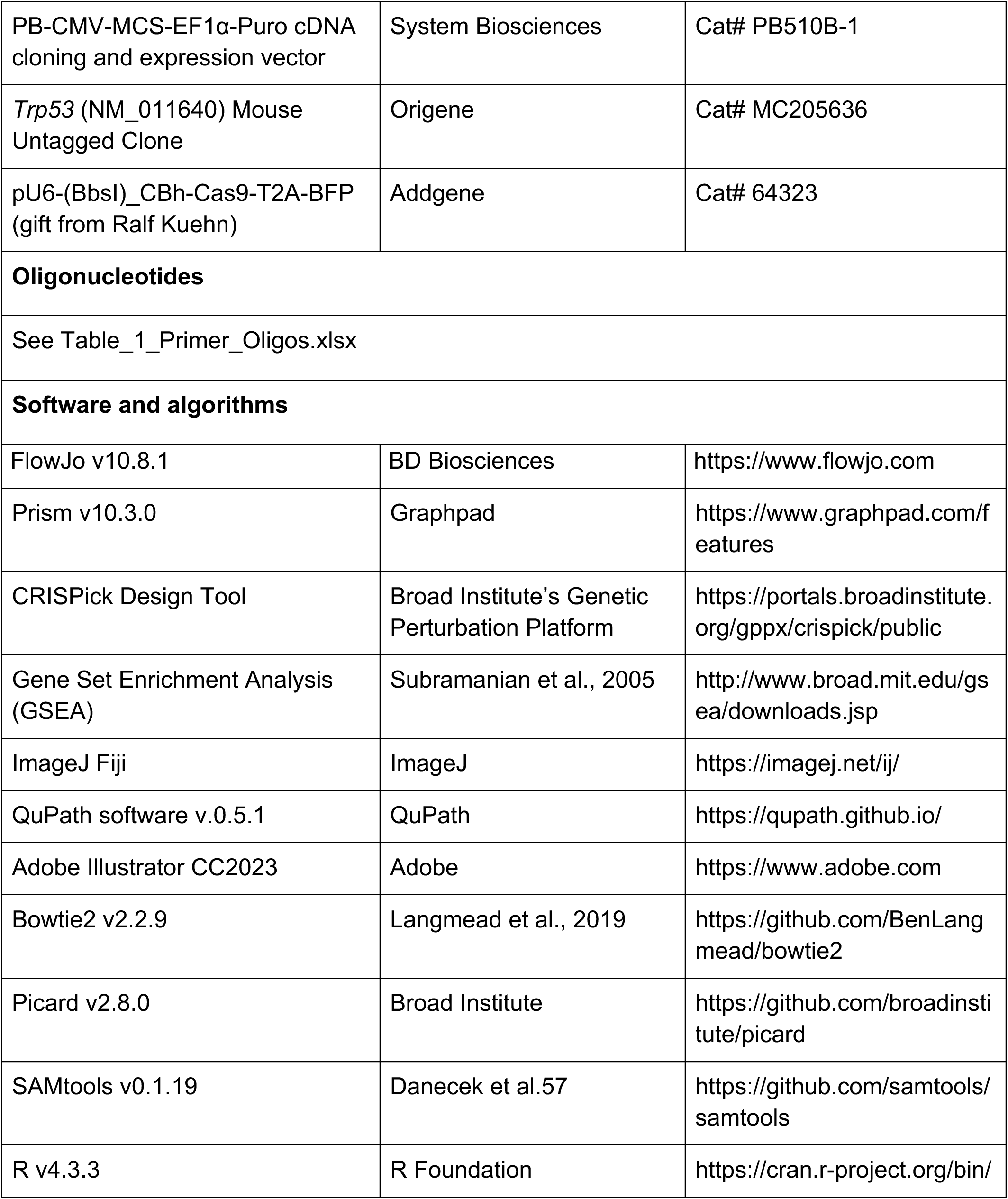
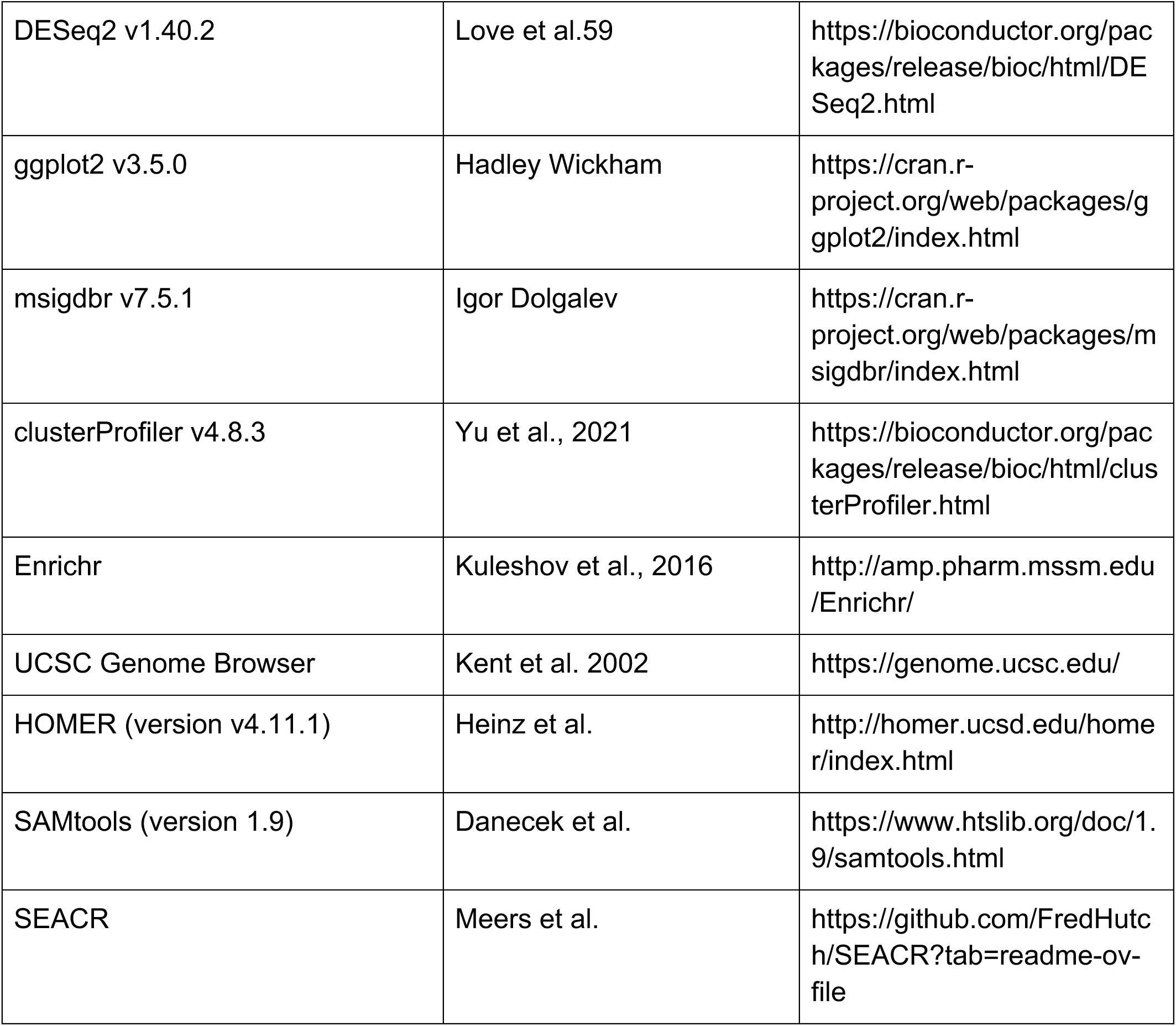

## RESOURCE AVAILABILITY

### Lead contact

Further information and requests for resources and reagents should be directed to the lead contact (P.A.S.).

### Materials availability

The lead contact can provide cell lines generated in this study upon request.

### Data and code availability

RNA-seq, PRO-seq, Exome-seq, and CUT&RUN data have been deposited in GEO and will be publicly available as of the date of publication. Accession numbers and secure tokens for reviewer access are listed in the key resources table.

RNA-seq - GSE275105. Secure token: ujstwuygpfsbpkz PRO-seq - GSE275106. Secure token: ytwtwwqkzvgjtgp Exome-seq - GSE275107. Secure token: sbcrgsaqndcplgx CUT&RUN - GSE275108. Secure token: ctcryaimjtefzav Table_1_Primer_Oligos.xlsx provides the primers and DNA oligos used to clone the guide RNA sequence in the Cas9 and gRNA-expressing plasmids.

The lead contact can provide any additional information required to reanalyze the data reported in this paper upon request.

## EXPERIMENTAL MODEL AND STUDY PARTICIPANT DETAILS

### Mice

We obtained C57Bl/6 mice from the Edwin L. Steele Laboratories, Massachusetts General Hospital, and The Jackson Laboratory (Bar Harbor, Maine). C57Bl/6 mice were bred and maintained in Cox-7 gnotobiotic animal facility at Edwin L Steele Laboratories. We housed the mice under specific pathogen-free conditions at the Koch Institute animal facility and the Massachusetts General Hospital animal facility. We gender-matched and age-matched the mice to be 6–10 weeks old at the time of experimentation. All experimental use of animals followed the Public Health Service Policy on Humane Care of Laboratory Animals and was approved by the Committee on Animal Care at MIT, the Institutional Animal Care and Use Committees (Massachusetts General Hospital/Harvard Medical School), and the Association for Assessment and Accreditation of Laboratory Animal Care International.

### PDAC cell lines

We received the KPC cells (*Kras^G12D/+^;Trp53^R172H/+^*) generated from the GEMMs of PDAC (*LSL- Kras^G12D/+^;LSL-Trp53^R172H/+^;Pdx-1-Cre*) from Dr. Ashok Saluja from the University of Miami. We received the HY19636 cells (*Kras^G12D/+^;Trp53^-/-^*) generated from the GEMMs of PDAC (*p48- Cre;Kras^LSL-G12D/+^;Trp53^loxP/+^* from Dr. Ronald DePinho, MD Anderson Cancer Center.

We cultured the PDAC cells in Dulbecco’s Modified Eagle Medium supplemented with 4.5 g/L D-glucose, L-glutamine, and 110 mg/L sodium pyruvate (Gibco), along with 10% heat-inactivated fetal bovine serum (Tissue Culture Biologics), on tissue culture-treated 10 cm plates (Corning). The cells were maintained at 37°C with 5% CO2 and passaged using 0.25% Trypsin-EDTA (Gibco) upon reaching ∼70% confluency. For tumor implantation, we trypsinized the cells and resuspended them in PBS at a concentration of 2.5 million cells/ml.

### Isogenic *Trp53^-/-^*, *Ccl2^-/-^*, *Cxcl1^-/-^*, *Cxcl5^-/-^*, and Cxcl1-enhancers-deleted (Δe8695, Δe8696, and Δe8697) PDAC cell lines

We generated CRISPR-deleted cells by first cloning specific sgRNAs into the Cas9 and Blue Fluorescent Protein (BFP) expressing pU6-(BbsI)_CBh-Cas9-T2A-BFP vector (Addgene). We designed the sgRNA sequences using the CRISPick Design Tool from the Broad Institute’s Genetic Perturbation Platform (see Table_1_Primer_Oligos.xlsx). The complementary top and bottom strands of the sgRNA oligos were synthesized with overhangs compatible with Type II restriction enzymes. We then phosphorylated the 5’ ends of these oligos with T4 Polynucleotide Kinase (NEB) at 37°C for 30 minutes.

Next, we employed the Golden Gate cloning method to ligate the annealed sgRNA cassettes into the BbsI-digested pU6-(BbsI)_CBh-Cas9-T2A-BFP vector. The reaction mixture consisted of the annealed sgRNA DNA cassette, the digested vector, 1X Tango Buffer (Thermo Scientific), 0.5 mM DTT, 0.5 mM ATP, FastDigest BbsI restriction enzyme (Thermo Scientific), and T7 DNA Ligase (NEB). We performed six cycles of digestion and ligation, alternating between 37°C for restriction digestion (5 minutes) and 21°C for ligation (5 minutes), for a total incubation time of 60 minutes. To eliminate excess unligated linear DNA, we treated the reaction with Plasmid-Safe ATP-Dependent DNase (Biosearch Technologies).

Following the ligation, we transformed the gRNA-cloned pU6-(BbsI)_CBh-Cas9-T2A-BFP plasmid into chemically competent Stbl3 E. coli cells (Thermo Scientific). We heat-shocked the cells at 42°C, incubated them on ice, and then recovered them in S.O.C. medium (Thermo Scientific) before plating them on LB agar containing Carbenicillin (Thermo Scientific). After overnight incubation, we selected single colonies, grew them in LB medium with Carbenicillin, and stored them as glycerol stocks at -80°C. Plasmid DNA was isolated from the selected colonies using the QIAprep Spin Miniprep Kit (Qiagen), and successful sgRNA insertion was confirmed by sequencing.

To achieve CRISPR-mediated gene deletion, we transfected 200 ng of the sequence-verified gRNA-cloned plasmid into 100,000 cells and seeded the previous day in a 24-well plate to reach 70-90% confluency. We prepared the transfection complexes by combining the purified plasmid DNA with Lipofectamine 2000 transfection reagent (ThermoFisher) in Opti-MEM I Reduced Serum Medium, incubated the mixture for 20 minutes, and then added it to the cells. Seventy-two hours post-transfection, we used FACS to select BFP-expressing single-cell clones, which we then isolated into 96-well plates. Finally, we sequenced these single-cell clones to confirm the presence of the intended gene edits - the deletion of the target region using a pair of gRNAs or the occurrence of frameshift or nonsense mutations in cases where a single gRNA was employed.

### Ectopic Expression of p53^R172H^ in *Trp53^-/-^* Cells

To test the sufficiency of p53^R172H^, we ectopically expressed *Trp53^R172H^* in *Trp53^-/-^* cells using a PiggyBac cDNA Cloning and Expression Vector. We PCR-amplified the *Trp53^R172H^* open reading frame (ORF) from the MC205636 plasmid using primers designed to add EcoRI and NotI restriction sites at both ends (see Table_1_Primer_Oligos.xlsx). The amplified product and the PiggyBac vector (System Biosciences) were digested with EcoRI and NotI at 37°C for 15 minutes. We then ran the digested products on a 1% agarose gel at 150 V for 2 hours and gel-purified the desired fragments: the vector backbone (6430 bp) and *Trp53^R172H^*insert (1819 bp). To prevent self-ligation, we dephosphorylated the vector using Alkaline Phosphatase (Thermo Scientific) and phosphorylated the restriction-digested *Trp53^R172H^* ORF with T4 Polynucleotide Kinase at 37°C for 30 minutes. We ligated the insert into the vector using T7 DNA Ligase at room temperature for 30 minutes.

The ligated plasmid construct was then transformed into One Shot TOP10 Chemically Competent E. coli cells (Thermo Scientific) via heat shock at 42°C, followed by recovery in SOC medium. After overnight incubation on LB agar plates containing Carbenicillin, we picked single colonies, cultured them in LB medium with Carbenicillin, and stored the cultures as glycerol stocks at -80°C. We isolated plasmid DNA from selected colonies using the Qiagen Miniprep kit and confirmed the successful insertion of the *Trp53^R172H^* ORF by sequencing.

For stable expression of *Trp53^R172H^* in *Trp53^-/-^*cells, we transfected 500 ng of the sequence-verified PiggyBac-*Trp53^R172H^*plasmid into one million *Trp53^-/-^* cells grown in a six-well plate, which had been seeded the previous day to achieve 60-80% confluency. We prepared the transfection complexes by combining the PiggyBac-*Trp53^R172H^*plasmid with PureFection (System Bioscience) in serum-free DMEM, vortexed the mixture for 30 seconds, incubated it for 15 minutes at room temperature and then added it dropwise to the cells. After allowing transposase activity for 72 hours, we applied puromycin selection. Although most cells died, a few colonies appeared after 10 days. We isolated individual colonies and verified the stable integration of the *Trp53^R172H^*construct. This process successfully generated *Trp53^-/-^* cells with stable, ectopic expression of the p53^R172H^ mutant protein.

To answer whether the Cxcl1 enhancer binding is specific to mutant p53, we tried expressing *Trp53^WT^* in *Trp53^-/-^* cells using multiple approaches, such as ectopic expression using piggyback plasmid, Dox-inducible system, or base-editing. Despite our efforts, we were unable to generate viable *Trp53^WT^*-expressing cells, consistent with previous findings that restoration of p53^WT^ induces senescence^87^, a therapeutic strategy under investigation in cancer clinical trials^77^.

## METHOD DETAILS

### Orthotopic pancreatic tumor implantation

We made a ∼1 cm incision in the skin and abdominal wall directly over the spleen to expose the pancreas. Using forceps, we gently spread the pancreas and injected 50,000 cells in a 20 µL volume into the pancreas, forming a bubble. We then closed the abdominal wall with absorbable sutures and used staples to close the skin. The staples were removed 7 to 10 days later to ensure proper skin healing and apposition.

### Tumor growth measurements and treatment

We measured tumor growth using high-frequency ultrasound. Every three days, we employed a Visualsonics Vevo 2100 ultrasound device equipped with a high-frequency 550S probe to noninvasively monitor tumor growth longitudinally under anesthesia. Once the tumors reached approximately 5 mm in diameter, we randomized the mice into different groups. Following randomization, we treated the mice with either IgG (control) or a cocktail consisting of 200 µg of α-PD-1 (BioXcell) and 100 µg of α-CTLA4 (BioXcell) in a 100 µl volume, administered every third day for a total of three doses.

### Tumor extraction and processing

We dissected tumors from the pancreas of mice, weighed them, and then minced and digested them in RPMI buffer containing 1 mg/ml Collagenase IV (Thermo Scientific), 1 mg/ml DNase I (Thermo Scientific), and 0.25 mg/ml Liberase (Sigma-Aldrich) at 37C for 20 minutes. We then strained the tumors through 100-micron filters using the back-end of a 1 ml syringe plunger into 50 ml conical Falcon tubes to create a single-cell suspension. We lysed red blood cells in 1 ml of ACK lysing buffer (Thermo Scientific) on ice for 2 minutes, followed by two washes with pre-chilled FACS buffer (PBS containing 1% FBS and 2 mM EDTA).

### Staining for flow cytometry

For intracellular protein staining, we added Brefeldin A at 1x to all reagents until the fixation/permeabilization step. We stained cells for 15 minutes on ice with Fixable Viability Dye eFluor 780 (eBioscience) or Fixable Viability Dye eFluor 506 (eBioscience) to differentiate between live and dead populations and ɑCD16/CD32 (BioLegend) to prevent non-specific antibody binding. After incubation, we washed the cells with FACS buffer and stained them for surface proteins using fluorophore-conjugated antibodies resuspended in FACS buffer at the specified dilutions for 20 minutes at 4°C. Following surface staining, we washed the cells twice with FACS buffer and fixed them using the Foxp3 Transcription Factor Fixation/Permeabilization buffer (eBioscience). Samples processed for myeloid cell staining and flow cytometry were not fixed. After fixation, we washed the cells twice with FACS buffer, then stained them for intracellular proteins in FACS buffer overnight at 4°C. To obtain the absolute cell count, we added Precision Count Beads (BioLegend) to the samples according to the manufacturer’s instructions. We washed the cells twice with FACS buffer before proceeding to acquiring data. We performed flow cytometry sample acquisition on an LSR Fortessa cytometer or FACSymphony A3 Cell Analyzer (BD Biosciences) and analyzed the collected data using FlowJo v10.8.1 software (BD Biosciences).

### Multiplex immunofluorescent staining of mouse tumors

We fixed tissues harvested from mice with 4% paraformaldehyde and embedded them in either paraffin or Tissue-Tek® O.C.T. Compound (Sakura Finetek). We sectioned the paraffin-embedded tissues at 4 µm and the O.C.T.-embedded tissues at 10 µm thickness. We deparaffinized and hydrated the paraffin-embedded slides, then retrieved epitopes using citrate-based Antigen Unmasking Solution (VECTOR Laboratories) at 98°C for 20 minutes. We blocked the slides with 5% normal donkey serum and 0.3% Triton-X in phosphate-buffered saline (PBS) for 1 hour at room temperature, followed by incubation overnight at 4°C with primary antibodies: 1:200 anti-CD8ɑ (Abcam), 1:250 anti-Neutrophil (Abcam), and 1:100 Arginase-1 (Cell Signaling Technology). After washing with PBS, we incubated the slides with a solution containing 4’,6- diamidino-2-phenylindole (DAPI) and secondary antibodies at a 1:200 dilution (711-166-152, Cy™3 AffiniPure™ F(ab’)₂ Fragment Donkey Anti-Rabbit IgG (H+L), Jackson ImmunoResearch; 712-606-150, Alexa Fluor® 647 AffiniPure™ F(ab’)₂ Fragment Donkey Anti-Rat IgG (H+L), Jackson ImmunoResearch) for 1 hour at room temperature, then mounted them with cover glass using VECTASHIELD Vibrance® Antifade Mounting Medium (VECTOR Laboratories). We imaged the stained slides at 20x magnification using a Zeiss Axio Scan Z1 (Carl Zeiss) and analyzed the captured image data to quantify CD8-positive cells or Gr-1 + Arginase-1-positive cells using QuPath software v.0.5.1^106^. For quantification, we randomly selected 5-8 regions of interest (ROIs) (500 x 500 µm) in each viable tumor tissue, calculated the average proportion and density of cells with the corresponding protein expression in every ROI, and reported these values as the representative value for each case.

### Cytokine and chemokine array

For the *Trp53^R172H/-^*and *Trp53^-/-^* cells, we plated 25,000 cells in 6-well tissue culture plates on day 1. To account for the background signal from the serum, we incubated media alone in an empty well. When the cells reached 10-20% confluency, we refreshed the media. After 48 hours, we collected the supernatants and centrifuged them to remove cellular debris. We then incubated the supernatants with the Mouse Cytokine Array C3 kit (RayBiotech) according to the manufacturer’s instructions. The membranes were imaged using the ChemiDoc MP Imaging System from Bio-Rad. We measured and analyzed cytokine intensities using the Protein Array Analyzer for Image J software, normalizing the results with control spots to determine relative cytokine intensities.

### ELISA

To measure the protein concentration of chemokine production across all cell lines, we coated 96-well ELISA plates (BioLegend) with 50 µl of 1 µg/ml anti-mouse CXCL1 (Leinco Technologies) or CCL2 (Bio-Techne) for the CCL2-specific ELISA. The coating was done overnight at 4^0^C in 50 mM Carbonate-Bicarbonate buffer (pH 9.5). After coating, we washed the plates three times with PBS containing 0.05% Tween 20 (wash solution) and blocked them with PBS containing 1% bovine serum albumin. We then prepared recombinant mouse CXCL1 protein (Bio-Techne) or recombinant mouse CCL2 protein (Bio-Techne) in a blocking buffer to generate a standard curve. Culture supernatants were centrifuged to remove cellular debris before being incubated on the pre-coated ELISA plates at room temperature for four hours, followed by thorough washing.

Next, we incubated the plates with 50 µl of 1 µg/ml anti-mouse CXCL1-biotin (Leinco Technologies) or anti-mouse CCL2-biotin (Bio-Techne) diluted in blocking buffer for two hours at room temperature, followed by additional washes. We then incubated the plates for 30 minutes with 50 µl of HRP Streptavidin (BioLegend) diluted at 1:1000 in blocking buffer at room temperature, followed by three washes. To detect the reaction, we added 50 µl of TMB Substrate Solution (BioLegend) for 4 minutes before stopping the reaction with 25 µl of Stop Solution (BioLegend). We analyzed the plates on an Infinite m200 microplate reader at a wavelength of 450 nm (Tecan). Both standards and culture supernatants were tested in duplicate.

We used commercially available ELISA kits for mouse CXCL5 (Thermo Scientific) and followed the manufacturer’s instructions to complete the assays.

### Quantitative real-time PCR (qRT-PCR)

We quantified the expression levels of specific genes using quantitative reverse transcription polymerase chain reaction (qRT-PCR). We extracted total RNA from *Kras^G12D/+^*;*Trp53^R172H/-^* and *Kras^G12D/+^*;*Trp53^-/-^*cells in tissue culture with the RNeasy Mini Kit (Qiagen), adhering to the manufacturer’s protocol. We determined the concentration and purity of the extracted RNA using Qubit Fluorometric Quantification (Thermo Scientific) and NanoDrop spectrophotometer (Thermo Scientific), respectively. We synthesized cDNA from 1 µg of total RNA with the High-Capacity cDNA Reverse Transcription Kit (Applied Biosystems), following the supplier’s instructions. We used the Power SYBR Green PCR Master Mix (Applied Biosystems) to perform qRT-PCR using a LightCycler® 480 System (Roche). Each reaction included 2 µL of cDNA template, 10 µL of SYBR Green master mix, and 0.5 µM of each forward and reverse primer (see Table_1_Primer_Oligos.xlsx), totaling 20 µL. The thermal cycling conditions were: initial denaturation at 95°C for 10 minutes, 40 cycles of 95°C for 15 seconds, and 60°C for 1 minute. We conducted a melting curve analysis to verify the specificity of the amplification products. We calculated relative gene expression normalizing to the housekeeping gene GAPDH. To ensure reproducibility and reliability, we performed all reactions in triplicate.

### RNA-seq

We extracted total RNA from *Kras^G12D/+^*;*Trp53^R172H/-^* cells, isogenic and syngenic *Kras^G12D/+^*;*Trp53^-/-^* cells, and *Trp53^-/^*^-^ + *pTrp53^R172H^* and *Trp53^-/-^* + *pEV* cells in tissue culture using the RNeasy Mini Kit (Qiagen) following the manufacturer’s protocol. We assessed RNA quality and concentration with an Agilent 2100 Bioanalyzer (Agilent Technologies). We prepared the library from RNA samples with an RNA integrity number (RIN) greater than 7.0 using the TruSeq Stranded mRNA Library Prep Kit (Illumina), according to the recommended protocol. Sequencing was performed on an Illumina NovaSeq 6000 platform, generating paired-end reads of 150 base pairs. We processed the raw sequence data for quality control using FastQC and trimmed adapters with Cutadapt. We aligned high-quality reads to the reference genome (mm10) using bowtie2 and performed transcript quantification using rsem. We conducted differential expression analysis with DESeq2 to identify significant changes in gene expression between conditions.

### Exome-seq

We performed Exome sequencing (Exome-seq) to confirm the mutations in the *Trp53* and *Kras* genes and to comprehensively analyze the coding regions of the genome. We extracted genomic DNA from the samples using a standard phenol-chloroform extraction method and quantified it with a Qubit fluorometer. Using the Agilent SureSelect mouse All Exon V5 kit, we captured the exonic regions according to the manufacturer’s protocol. We then subjected the captured libraries to paired-end sequencing on the Illumina NovaSeq 6000 platform. We processed the raw sequence data using the BWA-MEM algorithm to align it to the mouse reference genome (mm10). We performed subsequent variant calling using the GATK Best Practices pipeline and filtered the variants from BAM files using Mutect2. We annotated the identified variants with ANNOVAR and filtered them based on quality metrics, allele frequency, and predicted functional impact. Finally, we used CNVkit to determine the copy number status of the mutated positions from BAM files.

### PRO-seq

We adapted precision run-on sequencing (PRO-seq) from a previously published paper^107^. We prepared tissue culture cells for the nuclear run-on reaction by cell permeabilization. We removed the tissue culture medium, rinsed the cells with PBS, and placed the plates on ice. We then scraped the cells while still on ice, collected them into a 15 ml conical tube, and centrifuged them at 1,000g for 5 minutes. For nuclei isolation, we resuspended the pellet in ice-cold douncing buffer (10 mM Tris-Cl pH 7.4, 300 mM sucrose, 3 mM CaCl2, 2 mM MgCl2, 0.1% Triton X-100, 0.5 mM DTT, 0.1X Halt protease inhibitor, and 0.02 U/μl RNase inhibitor). We transferred it to a 7 ml dounce homogenizer (Wheaton, cat # 357542). After incubating on ice for 5 minutes, we dounced the cells 25 times with a tight pestle, transferred them back to the 15 ml conical tube, and centrifuged to pellet the nuclei. We washed the pellet twice in a douncing buffer. Then we resuspended the washed pellet in storage buffer (10 mM Tris-Cl pH 8.0, 5% glycerol, 5 mM MgCl2, 0.1 mM EDTA, 5 mM DTT, 1× Halt protease inhibitor, and 0.2 U/μl RNase inhibitor) at a concentration of 5 million nuclei per 50 μl of storage buffer. We flash-froze the suspension in liquid nitrogen and stored it at −80°C.

We mixed 10 × 10^6^ nuclei in 100 μl of storage buffer with 100 μl of 2x nuclear run-on buffer (10 mM Tris-HCl pH 8.0, 5 mM MgCl2, 1 mM DTT, 300 mM KCl, 1% Sarkosyl, 50 μM biotin-11- A/C/G/UTP, 0.2 units/μl RNase inhibitor) and incubated at 37°C for three minutes. We extracted total RNA, which includes biotin-labeled nascent RNA, using Trizol following the manufacturer’s protocol. We isolated and base-hydrolyzed the RNAs with a final concentration of 200 nM NaOH to an average size between 100-150 nucleotides. We isolated nascent RNAs with magnetic beads coated with streptavidin, followed by 3’ adapter ligation. After another round of biotin-streptavidin affinity purification, we removed the mRNA cap and ligated the 5’ adapter. Following the third biotin-streptavidin affinity purification, we generated cDNA by reverse transcription. We prepared PRO-seq libraries for sequencing using Illumina TruSeq small-RNA adaptors with twelve cycles of PCR.

To map the PRO-seq sequencing reads, we clipped adapters from PRO-seq reads using Cutadapt. We discarded reads shorter than 15 bp as they map to multiple sites in the genome. We mapped the reads to the mouse genome mm10 using bowtie2, requiring them to map uniquely to the genome and allowing up to 2 mismatches.

### Enhancer calling

We identified enhancers in *Kras^G12D/+^*;*Trp53^R172H/-^*cells as described previously^108^. We used dREG, a computational tool designed for genome-wide detection of regulatory elements from nascent transcription data, which leverages bidirectional transcription signals from PRO-seq to provide a high-resolution map of enhancers.

### CUT&RUN

We performed the Cleavage Under Targets and Release Using Nuclease (CUT&RUN) assay using the CUTANA™ ChIC/CUT&RUN Kit (EpiCypher, 14-1048) according to the manufacturer’s protocol. We prepared CUT&RUN with two different antibodies against p53, NF- κB subunit p65, p300, H3K4me3, and H3K27Ac. We prepared 500,000 cells per reaction and used concanavalin A-coated magnetic beads to capture cells. We added 0.5 ug antibody to permeabilized cells and incubated overnight at 4°C. We permeabilized the cells after overnight incubation with digitonin and introduced pAG-MNase to the cell-bead complexes to facilitate targeted chromatin digestion. We initiated the digestion by adding calcium chloride and incubating the samples at 4°C for 2 hours. To stop the reaction, we added EGTA and collected the supernatants containing the CUT&RUN-enriched chromatin. We purified the released DNA fragments using SPRIselect reagent and eluted the DNA. We quantified the eluted DNA using a Qubit fluorometer and prepared sequencing libraries using the NEBNext Ultra II DNA Library Prep Kit (New England Biolabs) according to the manufacturer’s instructions. The resulting libraries were sequenced on an Illumina NextSeq 500.

To map the CUT&RUN sequencing reads, we clipped adapters from CUT&RUN reads using Cutadapt. We discarded reads shorter than 15 bp as they map to multiple sites in the genome. We mapped the reads to the mouse genome mm10 in the --very-sensitive-local mode using bowtie2, requiring them to uniquely map to the genome. We then removed the PCR duplicates using the unique molecular identifiers (UMIs). We subsampled reads in the treatment group to match the reads in the control (IgG) group and called peaks using SEACR_1.3 (https://github.com/FredHutch/SEACR). We generated a union of p53^R172H^ peaks (n= 7,721) from two sets of peaks called with two p53 antibodies (Epicypher p53 antibody and Leica p53 antibody). As a control, we selected 100,000 random regions in the genome matched for p53^R172H^ CUT&RUN peak widths and filtered for expression levels of more than 1 CUT&RUN reads.

We used the Hypergeometric Optimization of Motif EnRichment (HOMER) software to identify enriched DNA sequences within CUT&RUN peaks. We extracted genomic sequences from these peaks and analyzed de novo motif enrichment using the HOMER motif discovery algorithm. As a background control, we generated 100,000 random genomic regions, each 1,000 nucleotides in length.

### TPCA-1 treatment

We dissolved 10 mg of TPCA-1 in 1.193 ml of DMSO to create a 30 mM stock solution. We plated cells in 15 cm tissue culture-treated plates and allowed them to adhere. 24 hours after plating them, we treated the cells with TPCA-1 (final concentration of 5 µM). As a control, we treated cells with 0.1% DMSO. We harvested cells 24 hours after the treatment and proceeded with either qRT-PCR or CUT&RUN.

---

**Figure S1.**
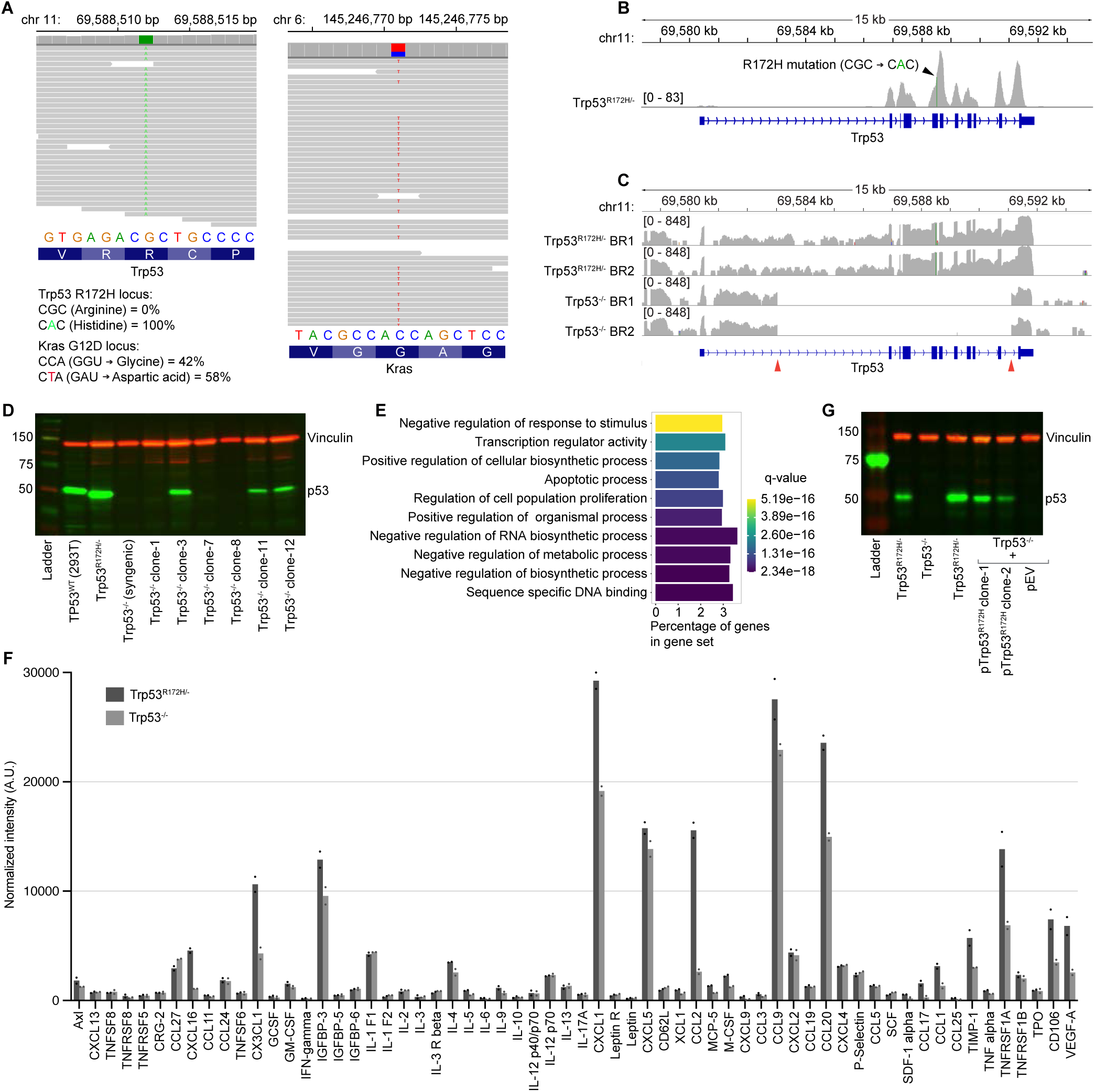
Genotype validation of isogenic PDAC cells and p53^R172H^-mediated gene expression changes. **A,** Exome-seq raw reads from *Trp53^R172H/-^* cells showing the R172H loci in the *Trp53* gene (left) and G12D loci in the *Kras* gene (right). Each gray horizontal line represents an Exome-seq read, and the colored nucleotide shows an alteration from the consensus sequence. The consensus DNA sequence and the corresponding amino acid sequences are shown at the bottom. **B**, An Integrative Genomics Viewer (IGV)^104^ screenshot of raw Exome-seq read pile-up at the *Trp53* locus. **C**, An IGV screenshot of RNA-seq reads in two biological replicates (BR) of *Trp53^R172H/-^*and *Trp53^-/-^* cells. The y-axis is in the log scale to show the depletion of RNA-seq reads in the deleted region of the *Trp53* gene (denoted by a red arrow). **D**, Western blot probed for p53 and Vinculin as the loading control in *Trp53^R172H/-^*and *Trp53^-/-^* single-cell clones. Clone-1, clone-7, and clone-8 are positive for *Trp53* deletion. Protein lysate from human embryonic kidney cell lines (293T), which contains wild-type p53, is also used as a control. The labels on the left indicate protein size in kilodaltons (kDa). **E**, Gene-ontology terms enriched in p53^R172H^-downregulated genes. **F**, Secreted chemokines and cytokines by the *Trp53^R172H/-^* and the isogenic *Trp53^-/-^* clone-1 cells quantified using a chemokine array. Bars represent the average of two measurements. **G**, Western blot probed for p53 and Vinculin as the loading control in single-cell clones of ectopically expressed p53^R172H^ in the *Trp53^-/-^* cells using a *Trp53^R172H^* cDNA expression cassette in piggyback vector (*Trp53^-/^*^-^ + *pTrp53^R172H^*). Clone-1 and clone-2 are positive for *Trp53^R172H^* insertion. The labels on the left indicate protein size in kilodaltons (kDa).

**Figure S2.**
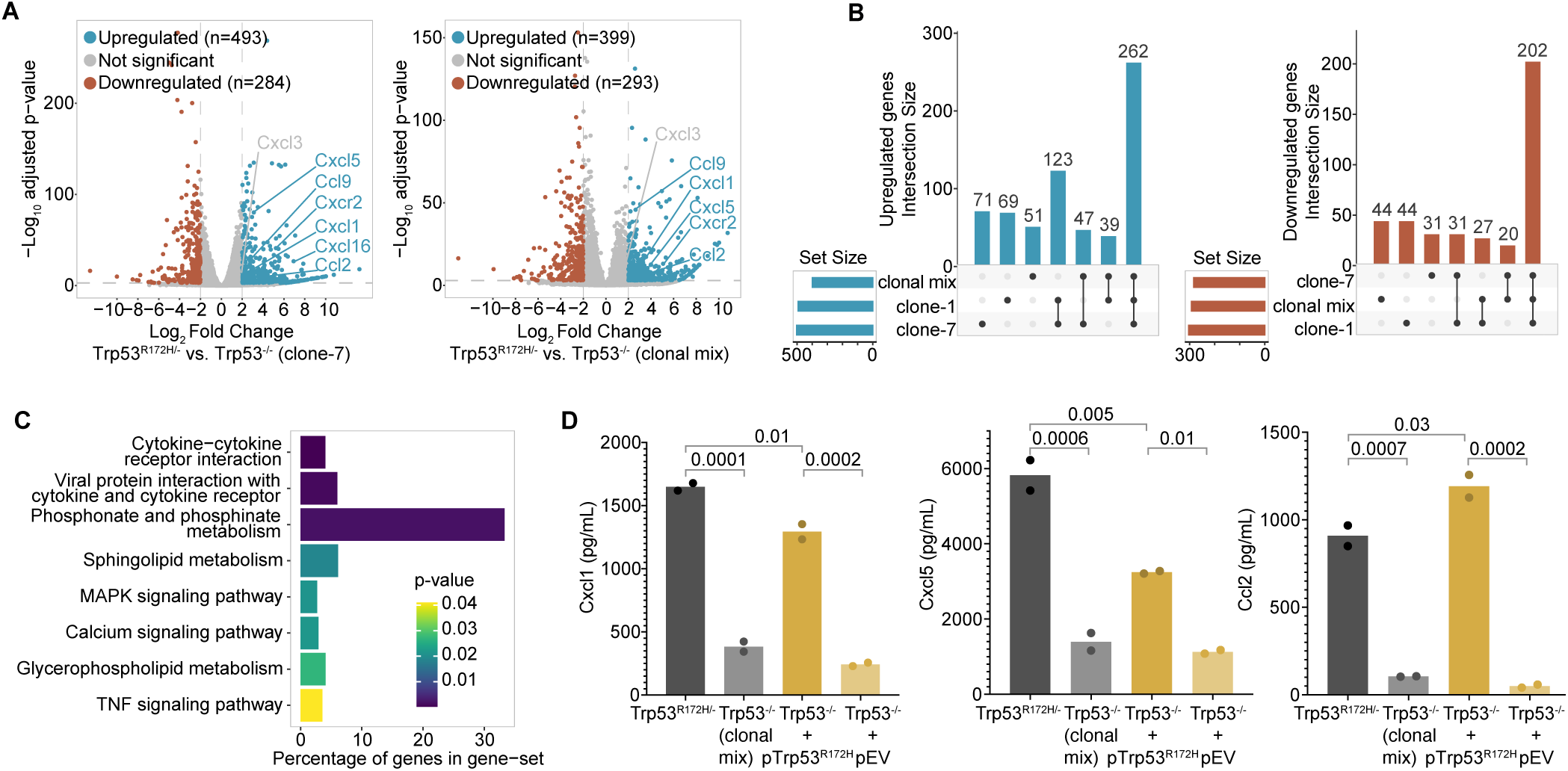
Validation of p53^R172H^-amplified expression of chemokine genes in additional single-cell clone and clonal mix population of *Trp53^-/-^* isogenic PDAC cells. **A,** Volcano plot of RNA-seq TPM showing the differentially expressed genes between *Trp53^R172H/-^* and a different single-cell clone (left) and clonal-mix (right) of *Trp53^-/-^* isogenic cells. Significantly upregulated and downregulated genes (adjusted p-value < 0.001 and four-fold change in normalized counts) are shown in blue and red, respectively, and the significantly upregulated chemokine genes are labeled in blue. **B,** Upset plot showing the overlap in significantly upregulated (left) and downregulated (right) genes in RNA-seq among two single-cell clones and a clonal-mix population of *Trp53^-/-^* cells. **C,** Gene sets enriched in p53^R172H^-upregulated genes (n = 262) common in the two single-cell clones and a clonal-mix population of *Trp53^-/-^* cells. Gene Ontology analysis was performed using Enrichr against the KEGG Pathway Database. **D,** Quantification of the three chemokine genes under the p53^R172H^ control, measured by ELISA in the tissue culture media of the four isogenic cells. The clonal-mix population of *Trp53^-/-^* cells was used. P-values are calculated from a one-way ANOVA test followed by a post hoc test with Benjamini-Hochberg correction.

**Figure S3.**
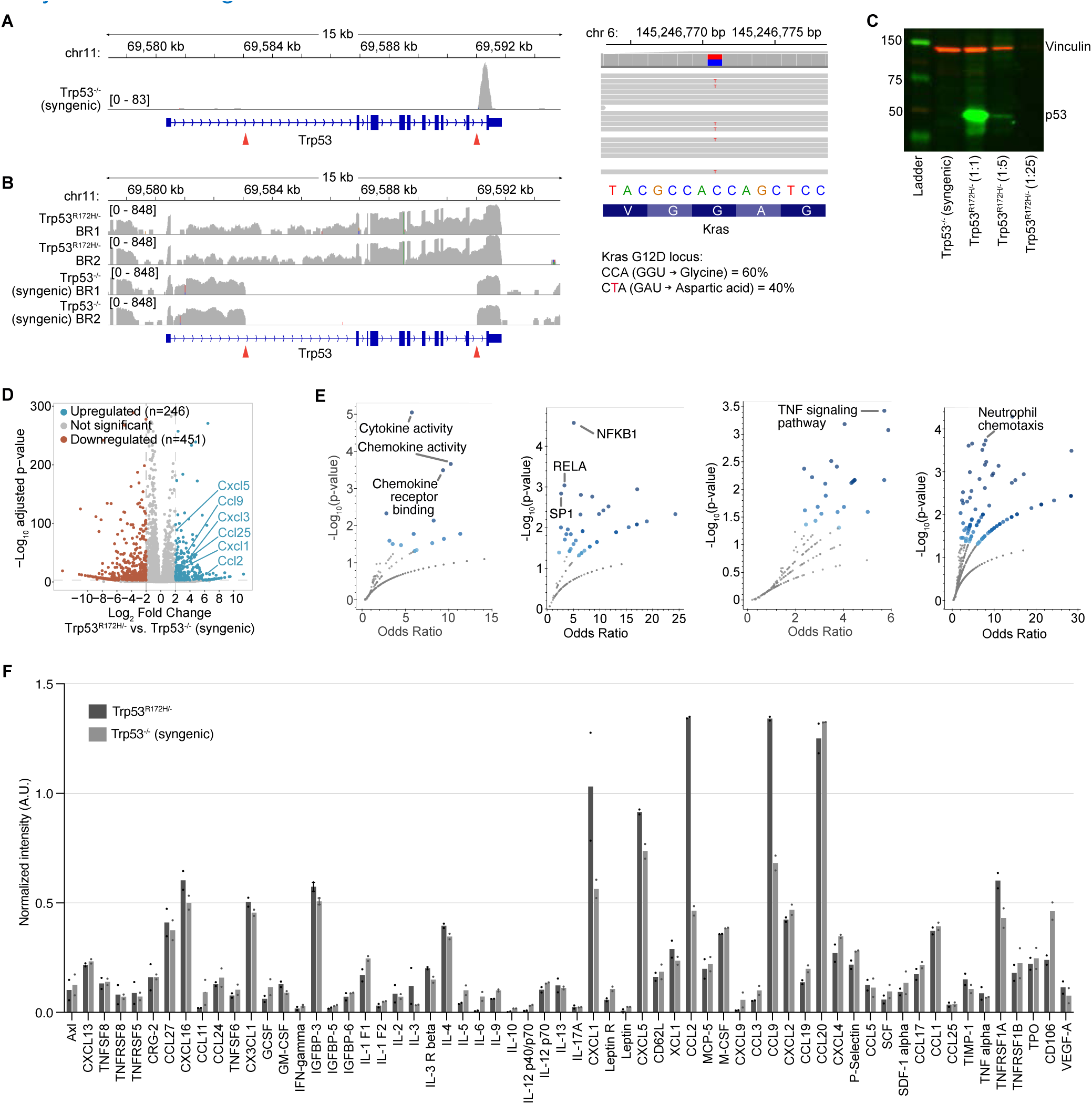
Genotype validation of syngeneic p53^NULL^ PDAC cells and p53^R172H^-mediated gene expression changes. **A,** An IGV screenshot of raw Exome-seq read pile-up at the *Trp53* locus of the *Trp53^-/-^* syngeneic cells (left). A red arrow denotes the deleted region of the *Trp53* gene. Exome-seq raw reads showing the G12D loci in the *Kras* gene of the *Trp53^-/-^* syngeneic cells (right). Each gray horizontal line represents an Exome-seq read, and the colored nucleotide shows an alteration from the consensus sequence. The consensus DNA sequence and the corresponding amino acid sequences are shown at the bottom. **B**, An IGV screenshot of raw RNA-seq reads in two biological replicates of *Trp53^R172H/-^*and *Trp53^-/-^* syngeneic cells. The y-axis is in the log scale to show the depletion of RNA-seq reads in the deleted region of the *Trp53* gene (denoted by a red arrow). **C,** Western blot probed for p53 (vinculin as the loading control) in the *Trp53^-/-^* syngeneic cells. A 5-fold serial dilution of protein lysate from the *Trp53^R172H/-^* cells is shown for the comparison of the p53 levels. The labels on the left indicate protein size in kilodaltons (kDa). **D**, Volcano plot of RNA-seq TPM showing the differentially expressed genes between the *Trp53^R172H/-^* and the *Trp53^-/-^* syngeneic cells. Significantly upregulated and downregulated genes (adjusted p-value < 0.001 and four-fold change in normalized counts) are shown in blue and red, respectively, and the significantly upregulated chemokine genes are labeled in blue. **E,** Pathways (left), transcription factor targets (central-left), signaling pathways (central-right), and immune compartment (right) enriched in p53^R172H^-upregulated genes. Blue dots represent significant gene sets (p-value < 0.05), and the darker color represents higher significance. **F**, Secreted chemokines and cytokines by the *Trp53^R172H/-^* and the syngeneic *Trp53^-/-^* cells quantified using a chemokine array. Bars represent the average of two measurements.

**Figure S4.**
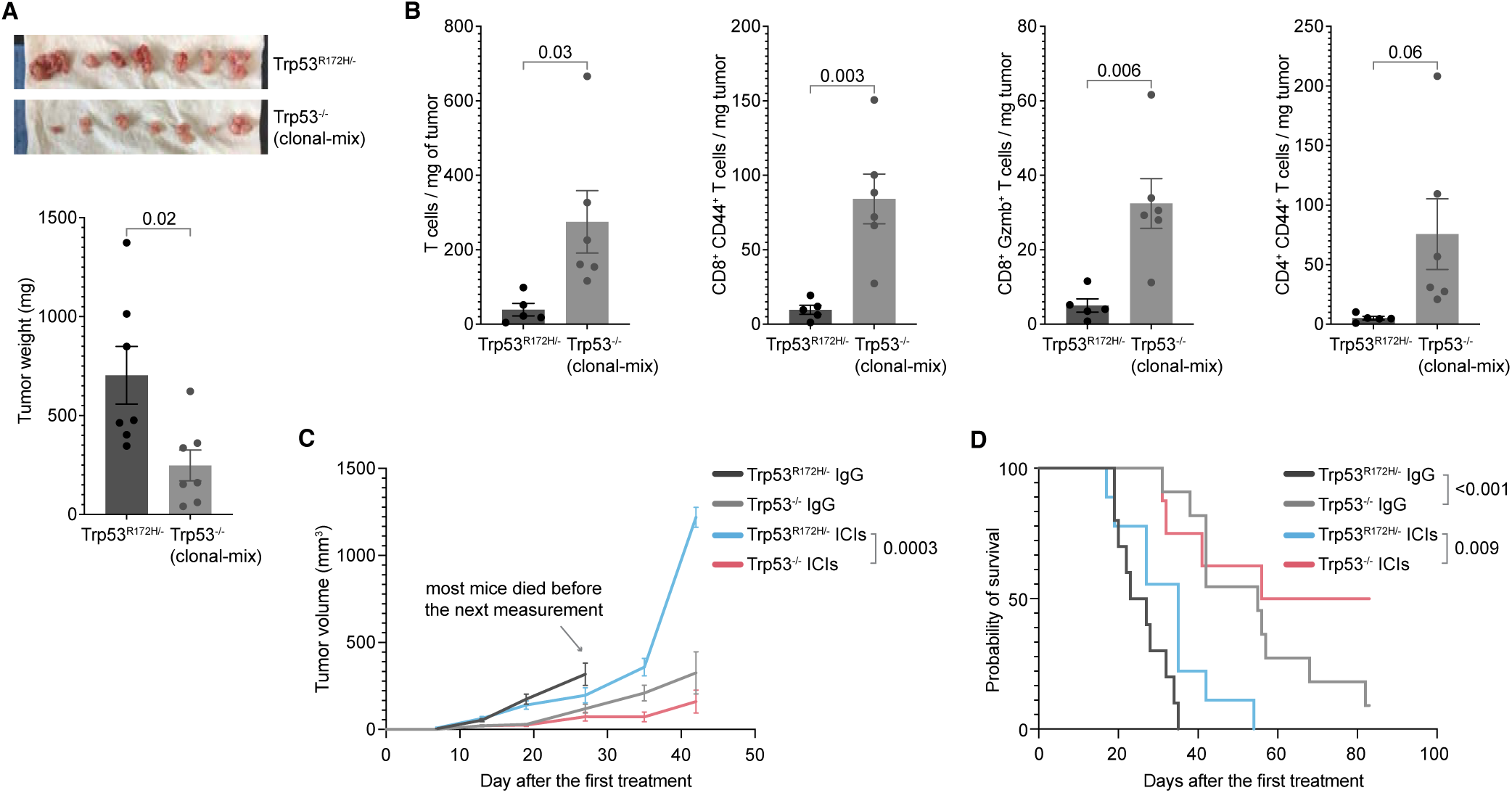
p53^R172H^-mediated immunosuppression and insulation from ICIs. **A,** Weights of the *Trp53^R172H/-^* and *Trp53^-/-^* clonal-mix tumors. The p-value is calculated using a two-tailed t-test. **B**, Effect of the *Trp53* status on activated and cytotoxic T cell infiltration in PDAC tumors. *Trp53^-/-^* clonal-mix cells were used. P-values are calculated using a two-tailed t-test. **C,** A replicate cohort (different from Figure 2F) of ICIs treatment showing the effect of the *Trp53* status and ICIs on PDAC tumor growth. Control mice were treated with IgG. Tumor volumes from the last measurement with at least three mice left in the cohort were used to calculate p- values using a two-tailed t-test. **D**, Kaplan-Meier survival curves in a replicate cohort of ICIs treatment showing the effect of *Trp53* status and ICIs on the survival of mice implanted with either *Trp53^R172H/-^* or *Trp53^-/-^* cells. Control mice were treated with IgG. P-values are calculated using a log-rank (Mantel-Cox) test.

**Figure S5.**
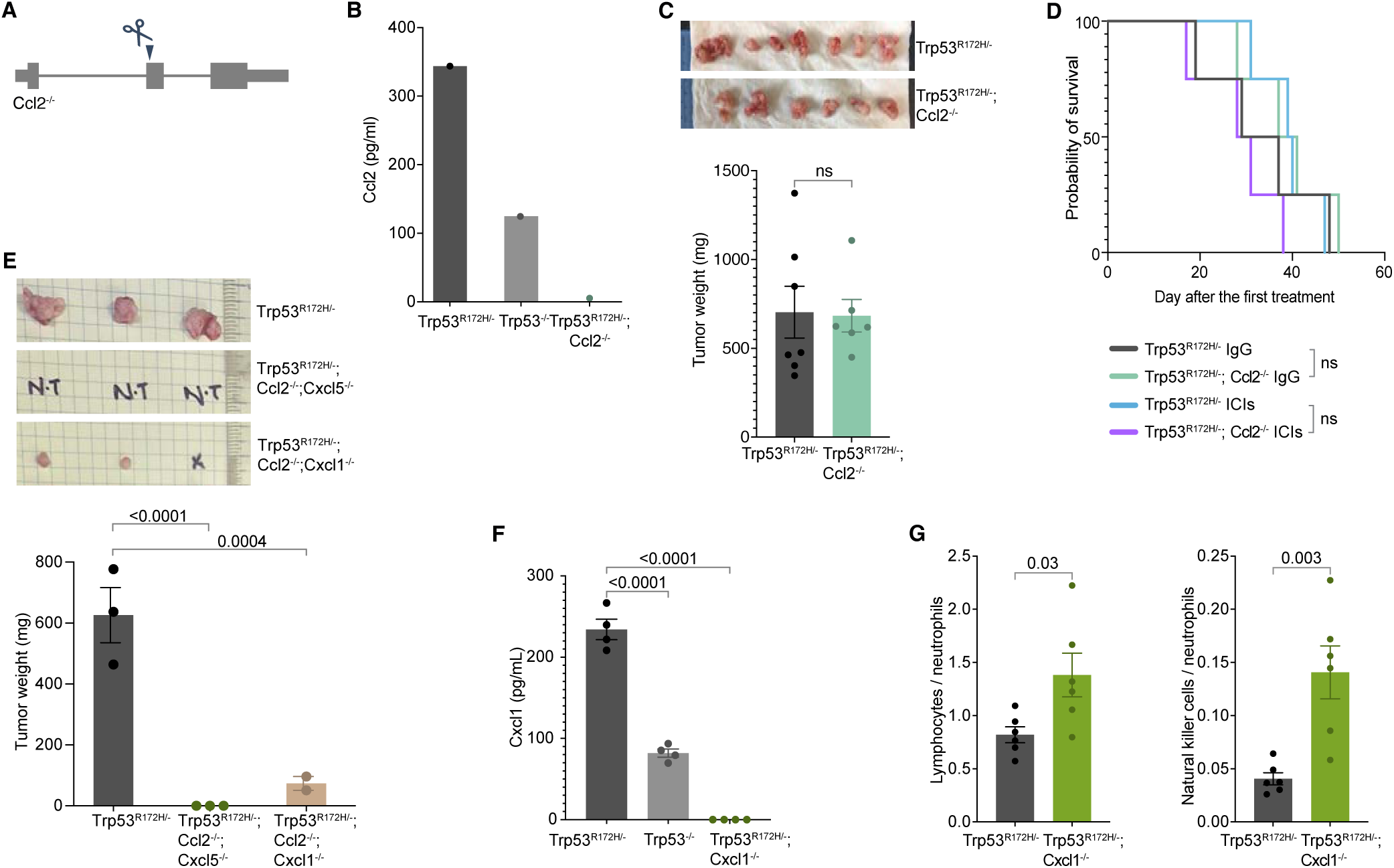
Ccl2 does not mediate the immunosuppressive role of p53^R172H^. **A,** Generation of *Trp53^R172H/-^*;*Ccl2^-/-^* isogenic cells from the parental *Trp53^R172H/-^* cells. A single guide-RNA-mediated genome editing using CRISPR/Cas9 resulted in a frameshift mutation. **B**, Quantification of the Ccl2 chemokine levels in selected *Trp53^R172H/-^*;*Ccl2^-/-^*single-cell clones and comparison with the *Trp53^R172H/-^* and *Trp53^-/-^* cells. **C**, Weights of the *Trp53^R172H/-^*;*Cxcl1^-/-^* tumors compared with the *Trp53^R172H/-^* tumors. P-values are calculated using a two-tailed t-test. **D**, Kaplan-Meier survival curves showing the effect of *Ccl2* status and ICIs in the survival of mice implanted with parental *Trp53^R172H/-^* or *Trp53^R172H/-^;Ccl2^-/-^*cells. Control mice were treated with IgG. Control mice were treated with IgG. P-values are calculated using a log-rank (Mantel-Cox) test. **E,** Weights of the *Trp53^R172H/-^*;*Ccl2^-/-^*;*Cxcl5^-/-^*and *Trp53^R172H/-^*;*Ccl2^-/-^*;*Cxcl1^-/-^*tumors compared with the *Trp53^R172H/-^* tumors in wild-type mice. P-values are calculated from a one-way ANOVA test followed by a post hoc test with Benjamini-Hochberg correction. **F,** Quantification of the Cxcl1 chemokine levels in *Trp53^R172H/-^*;*Cxcl1^-/-^*isogenic cells and comparison with the *Trp53^R172H/-^* and *Trp53^-/-^*cells. **G**, Effect of the *Cxcl1* status in the ratio of Lymphocytes and NK cells to Neutrophil in PDAC tumors.

**Figure S6.**
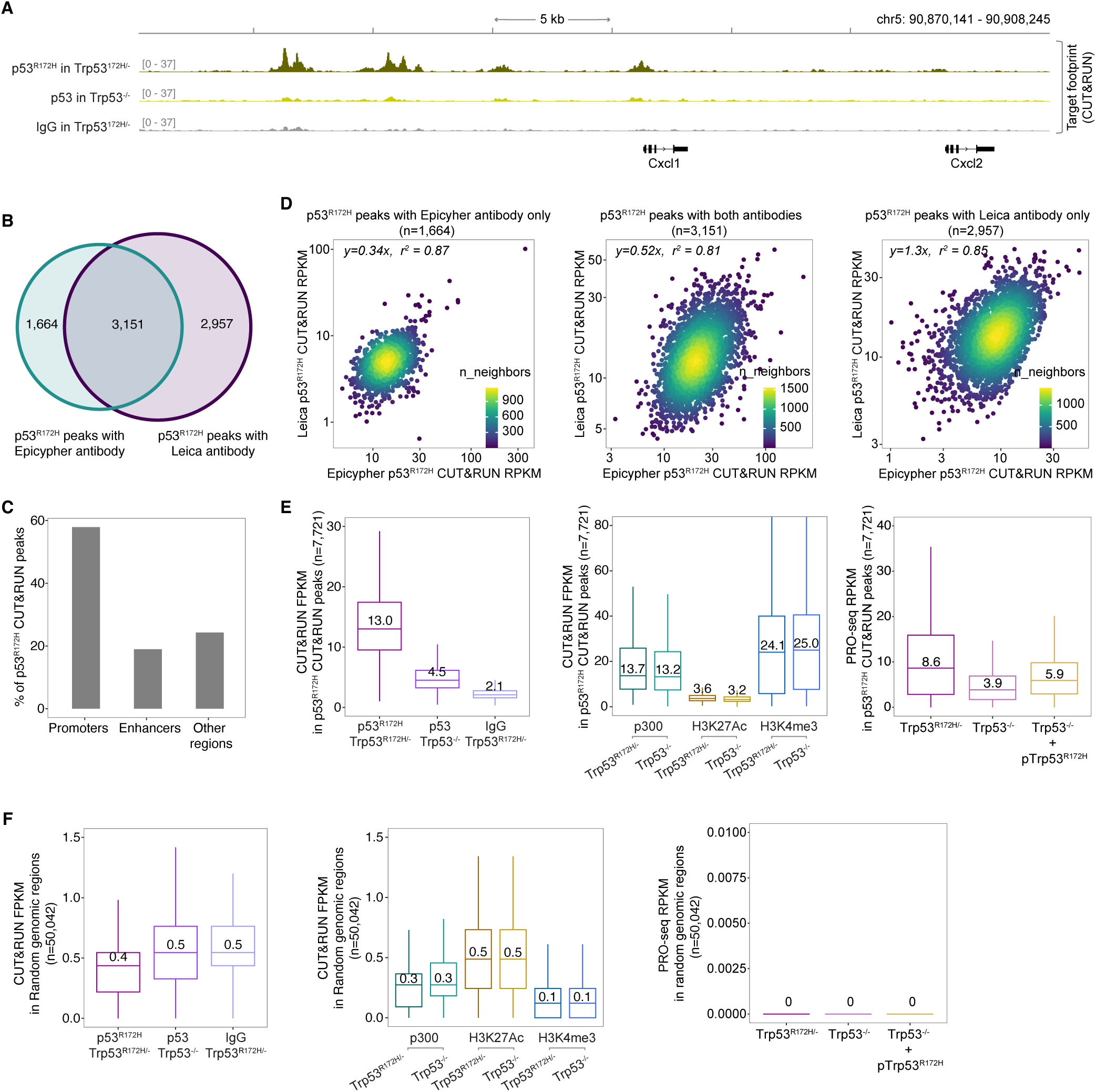
p53^R172H^ binding, but not the p300 and H3K27Ac levels, correlates with nascent transcription. **A,** p53^R172H^ occupancy at and around the *Cxcl1* gene in *Trp53^R172H/-^* and *Trp53^-/-^* cells using CUT&RUN with a different p53 antibody (see Methods). Non-specific IgG in *Trp53^R172H/-^* cells is used as a control. **B,** Overlap in p53^R172H^ CUT&RUN peaks with two different p53 antibodie. **C**, Distribution of p53^R172H^ CUT&RUN peaks in promoters (n=31,194), enhancers (n=11,893), and other regions, which may include intronic enhancers. **D,** Correlation between the CUT&RUN FPKM with Leica and Epicypher p53 antibodies in the three sets of unique and overlapping p53^R172H^ peaks. **E**, Quantification of p53^R172H^ occupancy (left), p300, and histone modification levels (middle), and nascent transcription (right) in the p53^R172H^ CUT&RUN peaks. **F**, Quantification of p53^R172H^ occupancy (left), p300, and histone modification levels (middle), and nascent transcription (right) in the random genomic regions (n=50,042, see Methods).

**Figure S7.**
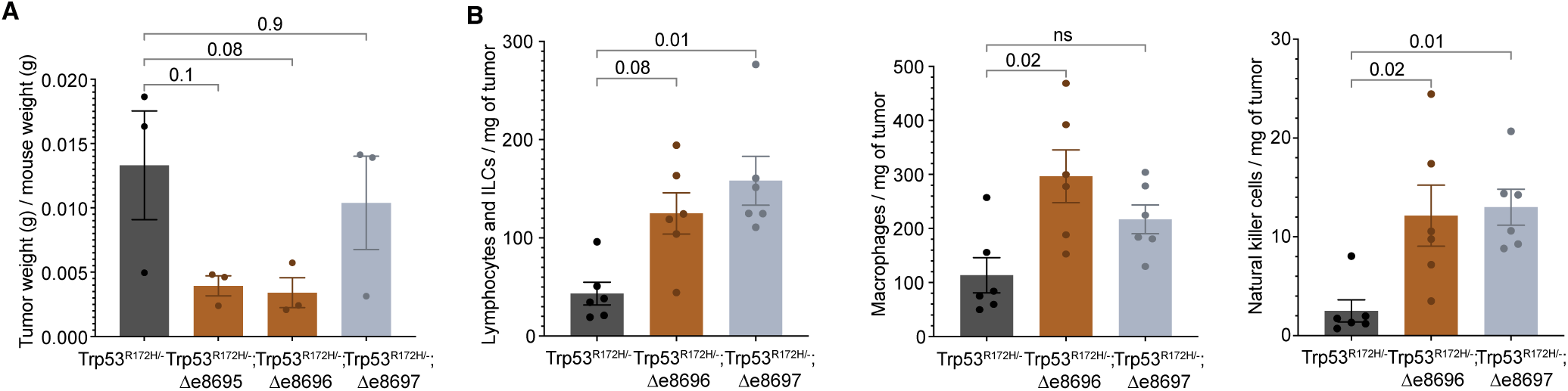
p53^R172H^-occupied Enhancers modulate PDAC TME. **A**, Weights from a replicate cohort of the enhancer-deleted isogenic tumors compared with the *Trp53^R172H/-^* tumors. P-values are calculated from a one-way ANOVA test followed by a post hoc test with Benjamini-Hochberg correction. **B,** Effect of the e8696 and e8697 status in lymphocytes and innate lymphoid cells, macrophage, and natural killer cell infiltration in PDAC tumors. P-values are calculated from a one-way ANOVA test followed by a post hoc test with Benjamini-Hochberg correction.

**Figure S8.**
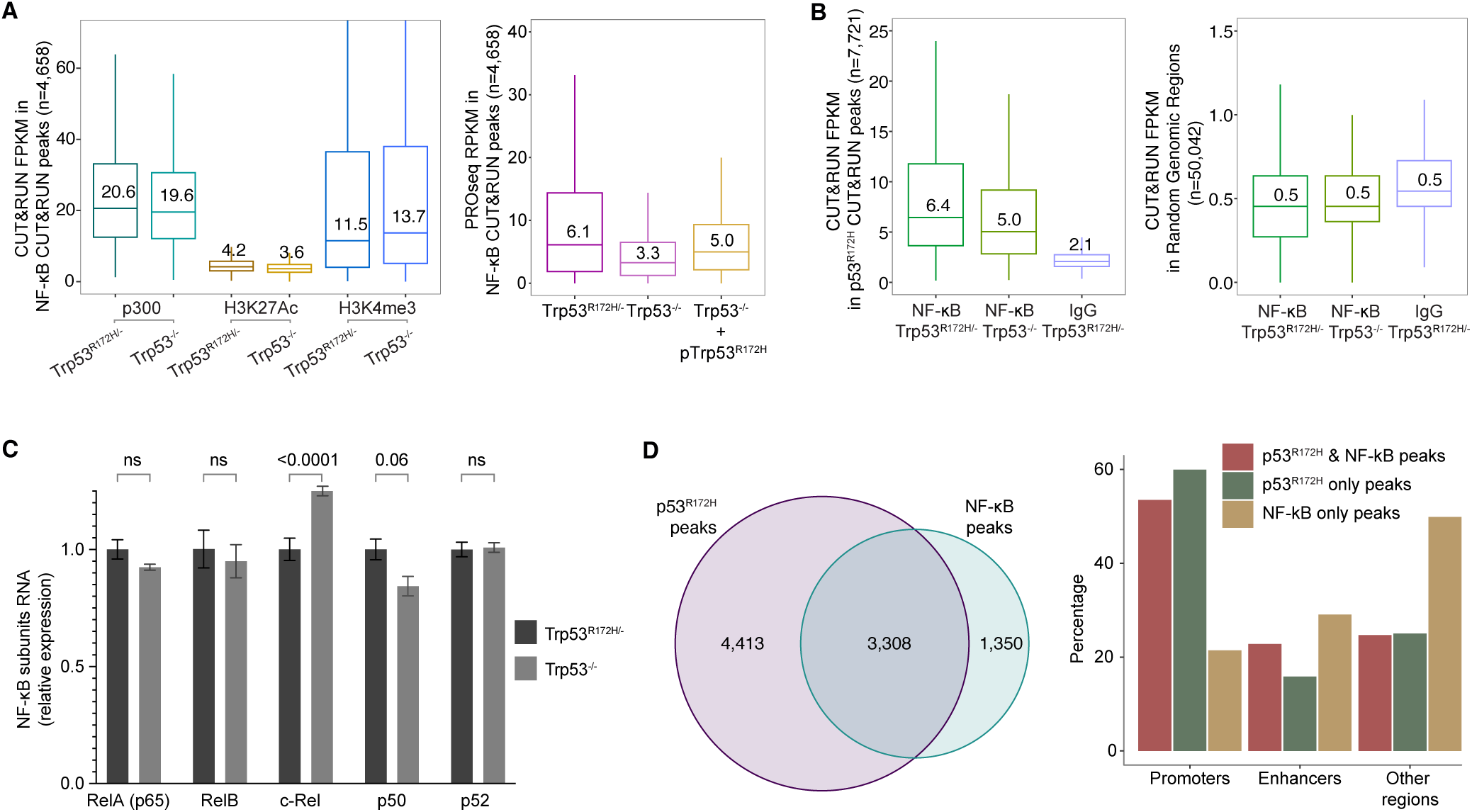
p53^R172H^ occupancy overlaps with NF-κB occupancy. **A,** Quantification of p300 and histone modification levels (left) and nascent transcription (right) in the NF-κB CUT&RUN peaks. **B**, Quantification of NF-κB occupancy in p53-occupied regions (left) and random genomic regions (right). **C**, Relative expression of NF-κB subunits in *Trp53^R172H/-^* and *Trp53^-/-^*cells quantified using RT- qPCR. **D**, Overlap in p53^R172H^ and NF-κB CUT&RUN peaks (left) and the distribution of p53^R172H^ and NF-κB CUT&RUN peaks in promoters (n=31,194), enhancers (n=11,893), and other regions, which may include intronic enhancers (right).

**Figure S9.**
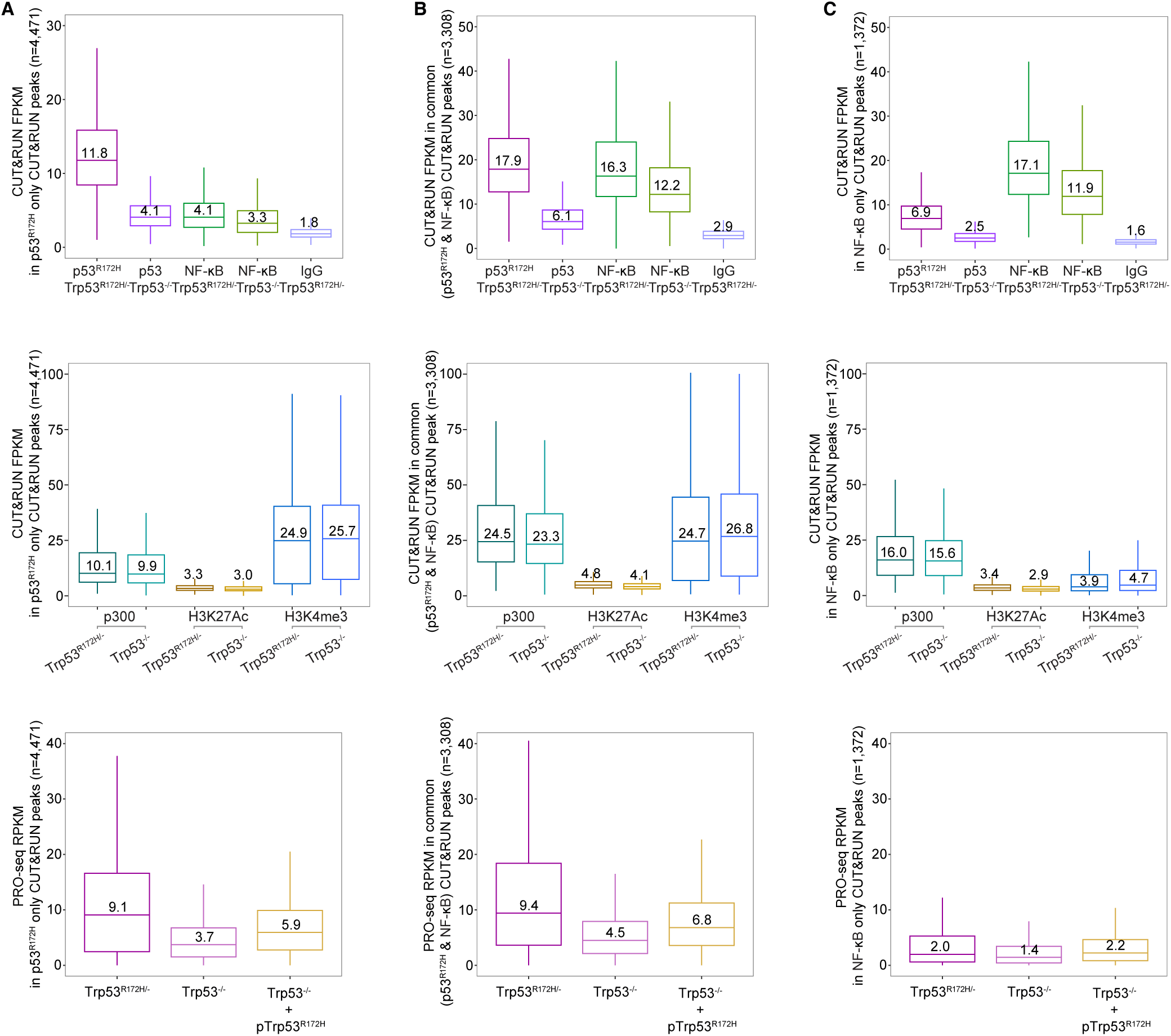
p53^R172H^ and NF-κB binding, but not the p300 and H3K27Ac levels, correlates with nascent transcription. **A,** Quantification of p53^R172H^ and NF-κB occupancy (top), p300 and histone modification levels (middle), and nascent transcription (bottom) in the unique p53^R172H^ peaks compared to the NF- κB peaks. **B,** Quantification of p53^R172H^ and NF-κB occupancy (top), p300 and histone modification levels (middle), and nascent transcription (bottom) in the common peaks in p53^R172H^ and NF-κB (n=3,308). **C,** Quantification of p53^R172H^ and NF-κB occupancy (top), p300 and histone modification levels (middle), and nascent transcription (bottom) in the unique NF-κB peaks compared to the p53^R172H^ peaks.

**Figure S10.**
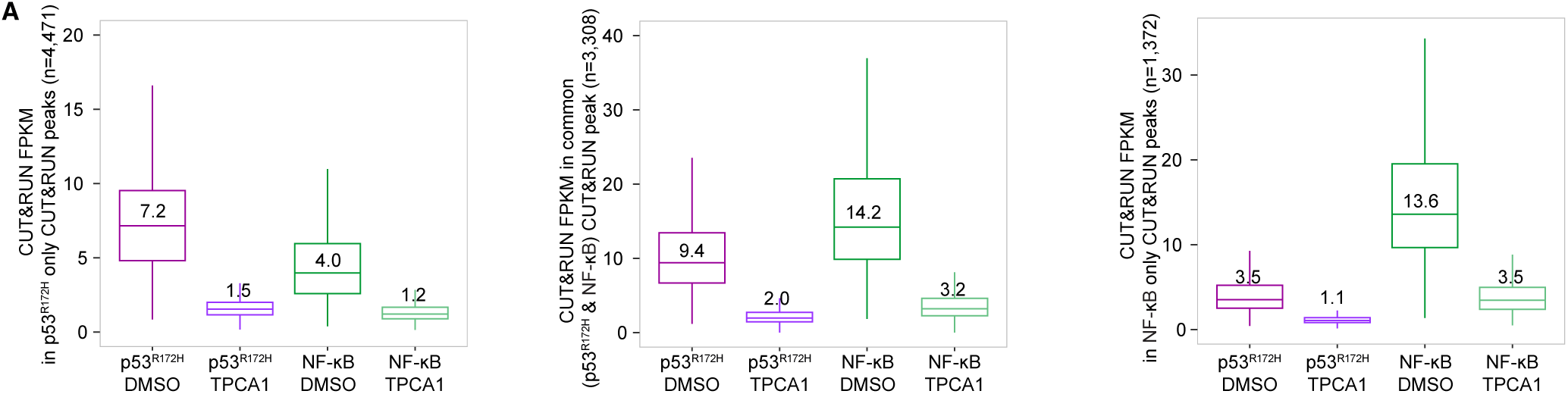
NF-κB inhibition abrogates p53^R172H^ and NF-κB occupancy. **A**, Effect of inhibiting NF-κB activation (5 uM TPCA-1) in NF-κB and p53^R172H^ occupancy in the unique p53^R172H^ peaks (left), the common p53^R172H^ and NF-κB peaks (middle), and the unique NF-κB peaks (right).

